# clampFISH 2.0 enables rapid, scalable amplified RNA detection *in situ*

**DOI:** 10.1101/2022.03.16.484659

**Authors:** Ian Dardani, Benjamin L. Emert, Yogesh Goyal, Connie L. Jiang, Amanpreet Kaur, Jasmine Lee, Sara H. Rouhanifard, Gretchen M. Alicea, Mitchell E. Fane, Min Xiao, Meenhard Herlyn, Ashani T. Weeraratna, Arjun Raj

**Affiliations:** Department of Bioengineering, School of Engineering and Applied Sciences, University of Pennsylvania, Philadelphia, PA, USA; Genomics and Computational Biology Graduate Group, Perelman School of Medicine, University of Pennsylvania, Philadelphia, PA, USA; Department of Cell and Developmental Biology, Feinberg School of Medicine, Northwestern University, Chicago, IL, USA; Center for Synthetic Biology, Northwestern University, Chicago, IL, USA; Genetics and Epigenetics, Cell and Molecular Biology Graduate Group, Perelman School of Medicine, University of Pennsylvania, Philadelphia, PA, USA; Department of Genetics, Perelman School of Medicine, University of Pennsylvania, Philadelphia, PA, USA; Department of Bioengineering, Northeastern University, Boston, MA, USA; Department of Biochemistry and Molecular Biology, Johns Hopkins School of Public Health, Baltimore, MD, USA; Department of Oncology, Sidney Kimmel Cancer Center, Johns Hopkins School of Medicine, Baltimore MD, USA; The Wistar Institute, Philadelphia, PA, USA

## Abstract

RNA labeling *in situ* has enormous potential to visualize transcripts and quantify their levels in single cells, but it remains challenging to produce high levels of signal while also enabling multiplexed detection of multiple RNA species simultaneously. Here, we describe clampFISH 2.0, a method that uses an inverted padlock design to efficiently detect and exponentially amplify signals from many RNA species at once, while also reducing time and cost compared to the prior clampFISH method. We leverage the increased throughput afforded by multiplexed signal amplification and sequential detection by demonstrating the ability to detect 10 different RNA species in over 1 million cells. We also show that clampFISH 2.0 works in tissue sections. We expect the advantages offered by clampFISH 2.0 will enable many applications in spatial transcriptomics.

## Introduction

Methods to label RNA in its cellular context have enabled us to visualize different aspects of gene expression. Initially, these approaches were dependent on an abundant and densely-packed target to generate sufficient signal (Singer and Ward 1982), though developments in probe synthesis and fluorescence microscopy enabled labeling and detection efficiencies sufficient to identify individual RNA molecules in fixed cells and tissue (Femino et al. 1998; Raj et al. 2008), a set of methods collectively termed single-molecule RNA fluorescence in situ hybridization (RNA FISH). These single-molecule RNA FISH methods localize multiple fluorescent dye molecules to a target RNA, typically using complementary DNA probes that, in early designs, were directly labeled with fluorescent dyes (Femino et al. 1998; Raj et al. 2008). This labeling approach, however, produces only weak fluorescent signals that hinders its use in high-background tissue sections and also requires long imaging times. To amplify the signal, there are now multiple single-molecule RNA FISH methods that build molecular scaffolds on the target RNA, providing a larger addressable sequence for fluorescent labeling. Each of these amplified methods, however, requires compromises in accuracy, multiplexing capacity, or cost. We recently described a method called clampFISH that is highly accurate with very high signal amplification, but was time-consuming, costly, and not validated for multiplexing beyond 3 targets. Here we present clampFISH 2.0, which solves these issues while retaining the strengths of the original method.

Amplification methods that build molecular scaffolds on their RNA do so with one of two approaches: using enzymes or using hybridization, each with their own drawbacks. Using multiple enzymes, rolling circle amplification generates a repeating DNA scaffold that can then be labeled fluorescently. Yet while there have been some reported improvements (Chen et al. 2018; Liu et al. 2021; Deng et al. 2017; Schneider and Meier 2017), enzymatic inefficiencies can leave many mRNA undetected (Lein, Borm, and Linnarsson 2017).

Alternatively, a number of methods forgo the use of enzymes, instead using hybridization-based approaches, including hybridization chain reaction (HCR) (Dirks and Pierce 2004b; Choi, Beck, and Pierce 2014; Choi et al. 2018), bDNA/RNAscope (Wang et al. 2012), FISH-STICs (Sinnamon and Czaplinski 2014), SABER-FISH (Kishi et al. 2019), and clampFISH (Rouhanifard et al. 2018). Although each of these hybridization-based methods offer various advantages, such as high signal gain or ease of repurposing for new targets, none can offer these features while offering rapid multiplexing beyond the 3-5 targets possible with spectrally distinguishable dyes.

Higher degrees of multiplexing beyond 3-5 targets requires the removal of the previous imaging cycle’s signal and iteratively re-probing a new set of targets (Lein, Borm, and Linnarsson 2017). The means by which this iterative procedure can be accomplished is constrained by the molecular scaffold design for each method.

With HCR, since the scaffold is built with dye-conjugated probes, higher degrees of multiplexing use DNAse digestion (Shah, Lubeck, Zhou, et al. 2016) of the previous cycle’s probes and repeat the entire protocol, thus substantially increasing the protocol time. Furthermore, the design of orthogonal HCR hairpin sets for multiplexing is complicated by the limited sequence diversity available for their short toehold sequences, such that to date, a maximum of 4-5 HCR amplifier probes have been used at once (Choi et al. 2018; Choi, Beck, and Pierce 2014). Unlike HCR, HiPlex RNAscope and non-branching SABER decouple the time-consuming amplification steps, which are done for all targets in parallel, and the short readout steps, in which fluorescently-labeled readout probes bind to a subset (eg. 3-5) of amplified scaffolds. After imaging in multiple fluorescent channels, the previous cycle’s signal is removed, and only the readout and imaging steps are repeated until all targets’ scaffolds have been probed. For higher degrees of multiplexing, this approach achieves an overall faster protocol, but it must also manage to remove the fluorescent signal without removing the remaining scaffolds. Thus, while the dye-coupled readout probes could otherwise be dissociated from their scaffolds with a stringent wash, this would also dissociate the scaffold components themselves, which are constructed via DNA hybrids. Hence, in lieu of a simple high-stringency wash, many published protocols call for photobleaching (Moffitt, Hao, Wang, et al. 2016) or fluorophore cleavage (Moffitt, Hao, Wang, et al. 2016; Xia et al. 2019) to eliminate the previous cycle’s signal. While non-branching SABER scaffolds (with up to ~10-fold amplification) can withstand the stringent wash to remove fluorescent readout probes, this approach has not been validated for the higher-gain branching SABER scaffolds (with up to ~450-fold amplification) (Kishi et al. 2019), likely because the branches’ shorter binding sequences could lead to their dissociation during the stringent washes. In contrast, clampFISH scaffolds could, in principle, withstand multiple stringent washes owing to their covalently locked configuration (Rouhanifard et al. 2018), although its other weaknesses (high probe cost, long protocol time) have limited the potential of this multiplexing approach.

Thus, there remains a need for an amplified RNA FISH method that permits accurate, high-gain, and flexible multiplexing. We reasoned that the original clampFISH (Rouhanifard et al. 2018), herein referred to as clampFISH 1.0, offered an excellent starting point for further development owing to a number of favorable attributes: its enzyme-free and step-wise amplification, its de-coupled amplification and readout, and its click chemistry-enabled boost in performance beyond the constraints imposed by oligonucleotide binding kinetics.

ClampFISH 1.0 probes are designed to form a looping scaffold off of a target nucleic acid molecule (Eg. RNA). Each probe begins as a linear DNA oligonucleotide which, upon hybridization to its target, forms a ‘C’ shape whose 5’ and 3’ ends are kept in close proximity via complementarity to a second sequence. Because the two ends are modified with alkyne (5’) and azide (3’) moieties, they can then be covalently linked using a click chemistry reaction (copper(I)-catalyzed azide–alkyne cycloaddition), which replaces the historically low-yield enzymatic ligation *in situ* (Lagunavicius et al. 2009). The click reaction circularizes the probe and, because of the helical structure of nucleic acid hybrids, stably loops it around its complementary strand (Rouhanifard et al. 2018).

The clampFISH 1.0 protocol begins by hybridizing multiple primary probes to the target, followed by multiple alternating amplification steps with secondary probes and tertiary probes. Successive rounds of amplification, in which 2 amplification probes can bind to each probe from the previous round, leads to a looping scaffold that grows exponentially in size. After every two steps, a click reaction is used to circularize the probes, thus stably linking the probes to one another. This amplified scaffold can then be labeled by hybridizing a fluorescent dye-coupled DNA readout probe to the many secondary or tertiary probes in the scaffold, thus localizing many fluorophores to the target to produce a single bright spot.

While clampFISH 1.0 offered a means for accurate, high-gain amplification, it suffered from a number of shortcomings that hindered its application. First, owing to multiple time-consuming amplification steps, clampFISH often took about 2.5 to 3 days to perform. Second, the method was plagued by many bright, mostly extracellular, non-specific spots that complicated downstream image analysis. Third, the high probe costs prohibited its widespread use, and these costs scaled poorly with additional gene targets, thus placing a practical limit on the method’s multiplexing potential. Lastly, it was unknown whether higher degrees of multiplexing, beyond spectral (3-gene) multiplexing, would be problematic, for example due to amplifier cross-reactivity or issues while removing readout probes.

Here, we introduce clampFISH 2.0, which addresses these shortcomings through multiple improvements to the original method, clampFISH 1.0. With an updated probe design and synthesis protocol, we greatly reduced the overall method cost, increased its scalability, and eliminated the extracellular non-specific spots. We optimized the method for faster amplification, reducing the pre-readout protocol time from ~2.5 days to about 18 hours, of which only 8 hours requires hands-on engagement, and we validated a 35 minute readout hybridization and stripping protocol for rapid 10-gene multiplexing. We show that clampFISH 2.0 is a fast, accurate, cheap, and flexible technology for multiplexed RNA detection *in situ*.

## Results

### Probe re-design and protocol optimization

ClampFISH 1.0’s primary probes were assembled with two gene-specific oligonucleotides that each required chemical modifications, substantially adding to the method’s cost. We therefore asked whether we could invert the primary probes’ orientation, such that the gene-specific RNA-binding oligonucleotide components could remain unmodified, and therefore cheaper, while incorporating the click chemistry modifications into a reusable, gene-independent oligonucleotide. In this scheme, we also add a separate ‘circularizer oligo’ to help ligate the primary probe, while keeping the orientation of the secondary and tertiary probes unchanged (Fig. 1a). This new primary probe design had the additional benefit of permitting larger-scale probe synthesis, since all of a gene’s primary probes could be ligated in a single pooled reaction (Fig. 1b). As a potential downside, the benefits of the new design could, in principle, come at the expense of specificity: the lack of a proximity ligation mediated by the target RNA molecule could allow for more non-specific probe self-ligation, because the proximity ligation is typically thought to increase specificity. Along with re-designing the primary probes, we also shortened the length of the secondary and tertiary probes (collectively referred to as ‘amplifier probes’), such that they can be made from a single commercially-produced oligonucleotide, thus simplifying their formerly 3-part synthesis.

**Figure 1:**
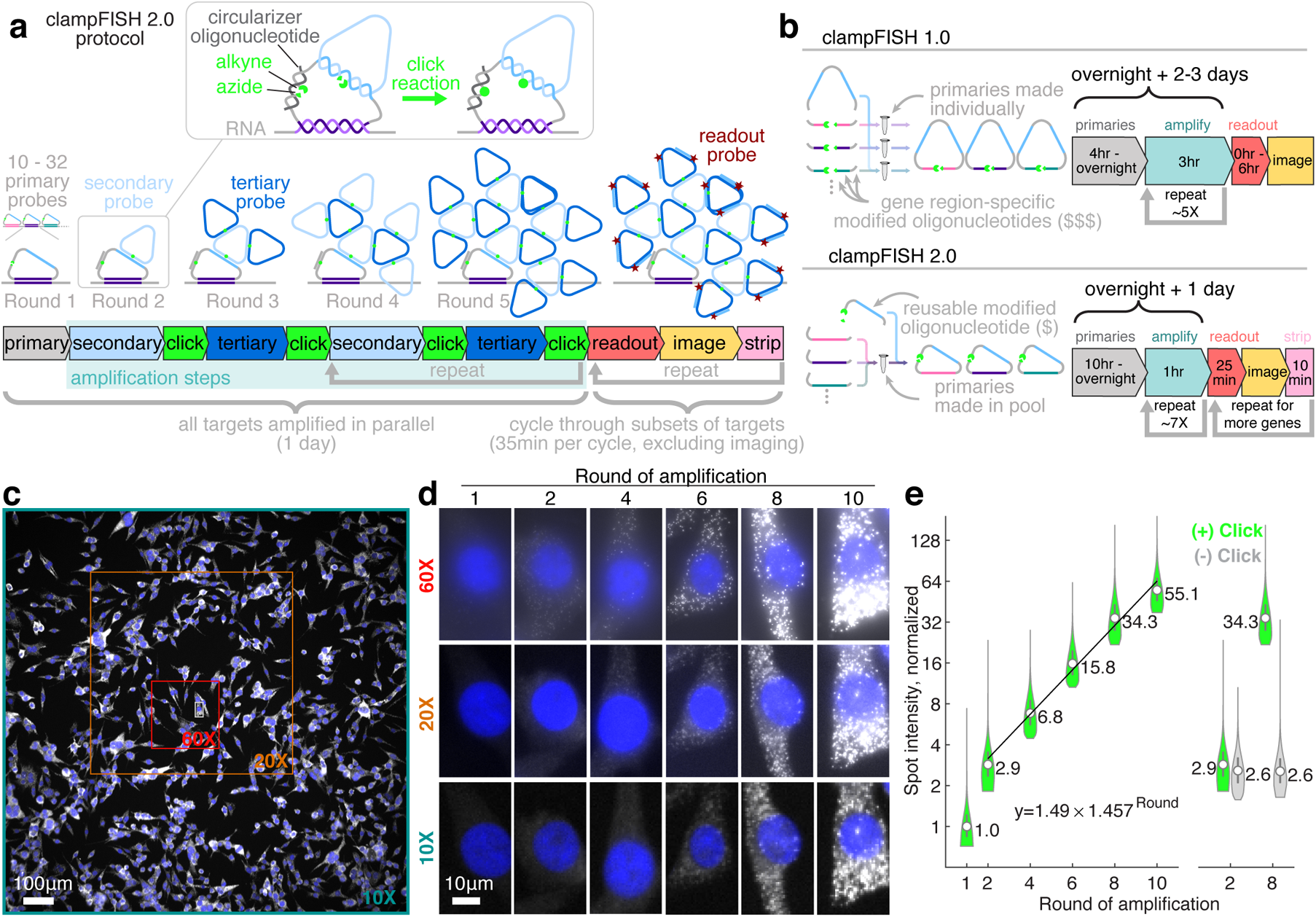
clampFISH 2.0 enables fast, cost-effective, exponential amplification of multiplexed RNA FISH signal in situ. **(a)** Schematic of clampFISH 2.0. **(b)** clampFISH 2.0 primary probes feature an inverted design, where oligonucleotides modified for use with click chemistry can be re-used for all probes in any primary probe set, rather than being designed specifically for one region on a particular gene, thus greatly reducing the overall probe cost from that of clampFISH 1.0. The new probe design also permits higher-throughput synthesis by allowing all primary probes for a given gene to be made in a pool. **(c)** *UBC* clampFISH 2.0 at round 10 in WM989 A6-G3 cells, imaged with a 10X objective, with the sizes of the smaller 20X and 60X fields of view overlaid. **(d)** UBC clampFISH 2.0 in WM989 A6-G3 cells shown at progressively higher rounds of amplification at 60X, 20X and 10X magnifications. **(e)** Left: *UBC* clampFISH 2.0 spots intensity (normalized to the median intensity from round 1) over progressively higher rounds of amplification, with the normalized median intensity from rounds 2, 4, 6, 8 and 10 fit to an exponential curve. Values shown are normalized median intensities. Right: spot intensities at rounds 2 and 8 when the copper catalyst is included (green) or not included (grey) in the click reaction. See Supplementary Figures 3 and 4 for data associated with additional targets and amplifier sets.

In addition to its high cost, the clampFISH 1.0 protocol was time-consuming, in large part because each round of amplification required approximately 3 hours. For example, the amplification protocol would require 2 days with 4-5 rounds of amplification, or 3 days for 6-8 rounds of amplification. To reduce the time of amplification, we sought to reduce the hybridization time by reducing secondary structure, as reported by (Gao, Wolf, and Georgiadis 2006; Zhang et al. 2014; Xia et al. 2019). With these new probe designs and additional optimization of the wash steps, click reaction, and buffer compositions, we reduced the time for a round of amplification from 3 hours to just 1 hour, which includes a 30 minute amplifier hybridization. This 3-fold speed improvement in amplification allows the full protocol, up to readout probe hybridization and imaging, to be performed with an overnight primary incubation (10 hr+) and about 8 hours the next day (Fig. 1b).

We wondered whether this updated scheme would still produce specific, amplified RNA FISH signal, as did the original clampFISH 1.0 method. We made primary probes for each of two separate mRNA targets (GFP mRNA, 10 probes; and *EGFR* mRNA, 30 probes) and tested their performance on a mixture of two cell lines known to express different RNAs: a WM989 A6-G3 H2B-GFP line, expressing the GFP sequence as mRNA, and a WM989 A6-G3 RC4 line grown in drug-containing media that we have shown to express high levels of *EGFR* mRNA (Shaffer et al. 2017; Emert et al. 2021; Goyal et al. 2021). We observed bright, amplified spots for the mRNAs specifically in the cells that were expected to express them (GFP spots in WM989 A6-G3 H2B-GFP cells, and *EGFR* spots in much greater numbers in WM989 A6-G3 RC4 cells; see Supplementary Figures 1 and 2), confirming the method’s specificity despite the new primary probes lacking an RNA-splinted proximity ligation. Furthermore, by introducing a number of centrifugation steps to the probe synthesis protocol, whereby we discard any material at the bottom of the tube, we largely eliminated the bright, non-specific spots seen with clampFISH 1.0 (Supplementary Figure 18)

We next sought to determine whether clampFISH 2.0 could exponentially amplify signal to a level that is detectable with lower-powered (20X/0.75NA and 10X/0.45NA) air objective lenses. We ran the clampFISH 2.0 protocol to varying stopping points: 1 round (primaries), 2 rounds (primaries and secondaries), 4 rounds (primaries, secondaries, tertiaries, and secondaries again), 6 rounds, 8 rounds, and 10 rounds, and hybridized readout probes to these scaffolds. Using low-powered magnification with large fields of view, we can reliably detect spots after amplification, thus demonstrating clampFISH 2.0’s capacity for high-throughput RNA detection (Fig. 1c,d). Furthermore, we observed an exponential rate of amplification, measured to be 1.406 to 1.678-fold per round (Fig. 1e, Supplementary Figure 3), implying the amplifier binding efficiency is 70 - 84% of the theoretical doubling of intensity per round. This exponential rate of growth did not appreciably slow down, even at the maximum number of rounds tested (round 10) where we detected up to ~100-fold signal amplification (Supplementary Figure 3), suggesting that an even brighter signal could be achievable with additional amplification. When we removed the copper catalyst from the click reaction, we saw no amplification from round 2 to round 8 (Fig. 1e, Supplementary Figure 4), suggesting that the scaffold’s growth requires looping of circularized amplifier probes to one another.

In order to achieve a higher degree of multiplexing, we needed to have a number of orthogonal sets of amplifier probes that had high gain and low off-target activity. We thus screened 15 amplifier probe sets, each used with primary probes targeting GFP mRNA or *EGFR* mRNA. Of these, we chose 10 sets of amplifier probes (1, 3, 5, 6, 7, 9, 10, 12, 14, and 15) with high gain and low off-target activity (amplifier set 11 was excluded based on its high number of off-target spots). We observed that an amplifier probe set’s gain for one RNA target strongly correlated with its gain on the other RNA target, suggesting that amplifiers can be used in a modular fashion with any set of primary probes without substantial primary-probe-specific effects on performance (Supplementary Figures 5 and 6). We also confirmed that amplifier probes do not cross-react with one another by showing that the spot intensities were equivalent when amplifier sets were used individually versus when they were used in a pooled mixture (Supplementary Figure 7) (we did not explicitly test whether amplifier probes interact with primary probes for which they were not designed).

Given the method’s capacity for fast, flexible multiplexed RNA detection, we next characterized its quantitative accuracy when used at low magnification, a capability useful for high-throughput imaging of large sample areas. We performed clampFISH 2.0 to round 8 (one round of primary probes and 7 rounds of amplifier probes) targeting three mRNAs (*EGFR*, *AXL*, and *DDX58*) with a range of expression levels. After the clampFISH 2.0 protocol, we hybridized conventional, unamplified single-molecule RNA FISH probes as a gold standard (Raj et al. 2008), which were designed to bind to non-overlapping sites on the same mRNA. We were able to observe many of the same spots with clampFISH 2.0 at 20X magnification that we saw using conventional single-molecule RNA FISH at 60X high magnification (60X), confirming the method’s high sensitivity and specificity (Fig. 2a). In addition to the 9X larger field of view and greater depth of field offered by 20X magnification compared with 60X magnification, we detected clampFISH 2.0 spots at 20X using shorter exposure times (100 milliseconds for *EGFR*, 250 milliseconds for *AXL*, and 500 milliseconds for *DDX58*) in comparison to the 2 second exposure time used with conventional single-molecule RNA FISH. Comparing the spot counts for multiple targets between clampFISH 2.0 at 20X magnification and conventional single-molecule RNA FISH at 60X magnification, we observed a high correlation, demonstrating that clampFISH 2.0 can be used as a higher-throughput replacement for conventional single-molecule RNA FISH (Fig. 2b). Even for a target (*DDX58*) expressed at low levels in a subset of cells, we were able to accurately identify cells with 3 or more RNAs (41-53% sensitivity, 97-99% specificity), thus supporting the ability of clampFISH 2.0 to reliably quantify even lowly-expressed genes at 20X magnification. Comparing spot counts from clampFISH 2.0 at 10X magnification to conventional single-molecule RNA FISH at 60X magnification, we saw a reduction in the correlation strength (eg. when targeting *AXL*, we observed an R^2^ of 0.740 to 0.773 at 10X magnification, versus an R^2^ of 0.891 to 0.899 for 20X magnification; see Fig. 2b and Supplementary Figure 8), suggesting that more accurate quantification using 10X magnification may require additional rounds of amplification beyond round 8. While the majority of clampFISH 2.0 spots lie in the cytoplasm, as expected, we also detected spots in the nucleus (Fig. 3b), a feature of clampFISH 2.0 that enables high-throughput analyses involving RNA localization or the rate of transcription.

**Figure 2:**
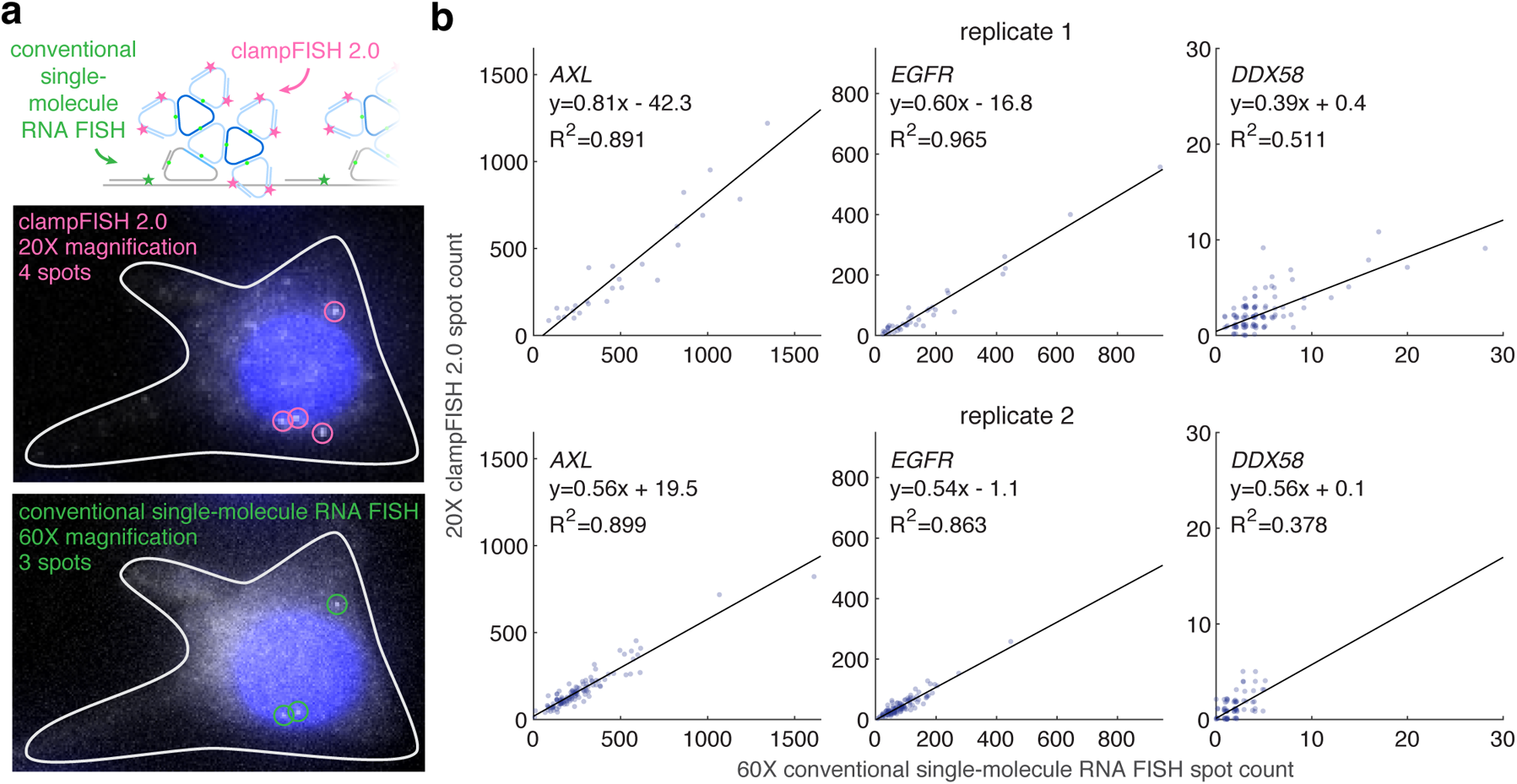
clampFISH 2.0 accurately quantifies RNA spot counts at low-powered magnification. **(a)** Top: Schematic depiction of labeling the same RNA with clampFISH 2.0 and conventional single-molecule RNA FISH, probing non-overlapping regions of the RNA. Middle: Image of *DDX58* clampFISH 2.0 spots with readout probes labeled in Alexa Fluor 594 and imaged at 20X magnification. Bottom: Image of conventional single-molecule RNA FISH (labeled with Cy3) targeting non-overlapping regions of *DDX58* at 60X magnification in the same cell. **(b)** We performed clampFISH 2.0 for 10 genes, amplified the 10 scaffolds in parallel to round 8, then added a single pair of readout probes to label a scaffold corresponding to *AXL* (left; in drug-resistant WM989 A6-G3 RC4 cells), *EGFR* (middle; in drug-resistant WM989 A6-G3 RC4 cells), or *DDX58* (right; in drug-naive WM989 A6-G3 cells). In two biological replicates (top: replicate 1; bottom: replicate 2), we counted spots for clampFISH 2.0 at 20X magnification (y-axis) and conventional single-molecule RNA FISH at 60X magnification (x-axis), which targeted non-overlapping regions of the same RNAs.

**Figure 3:**
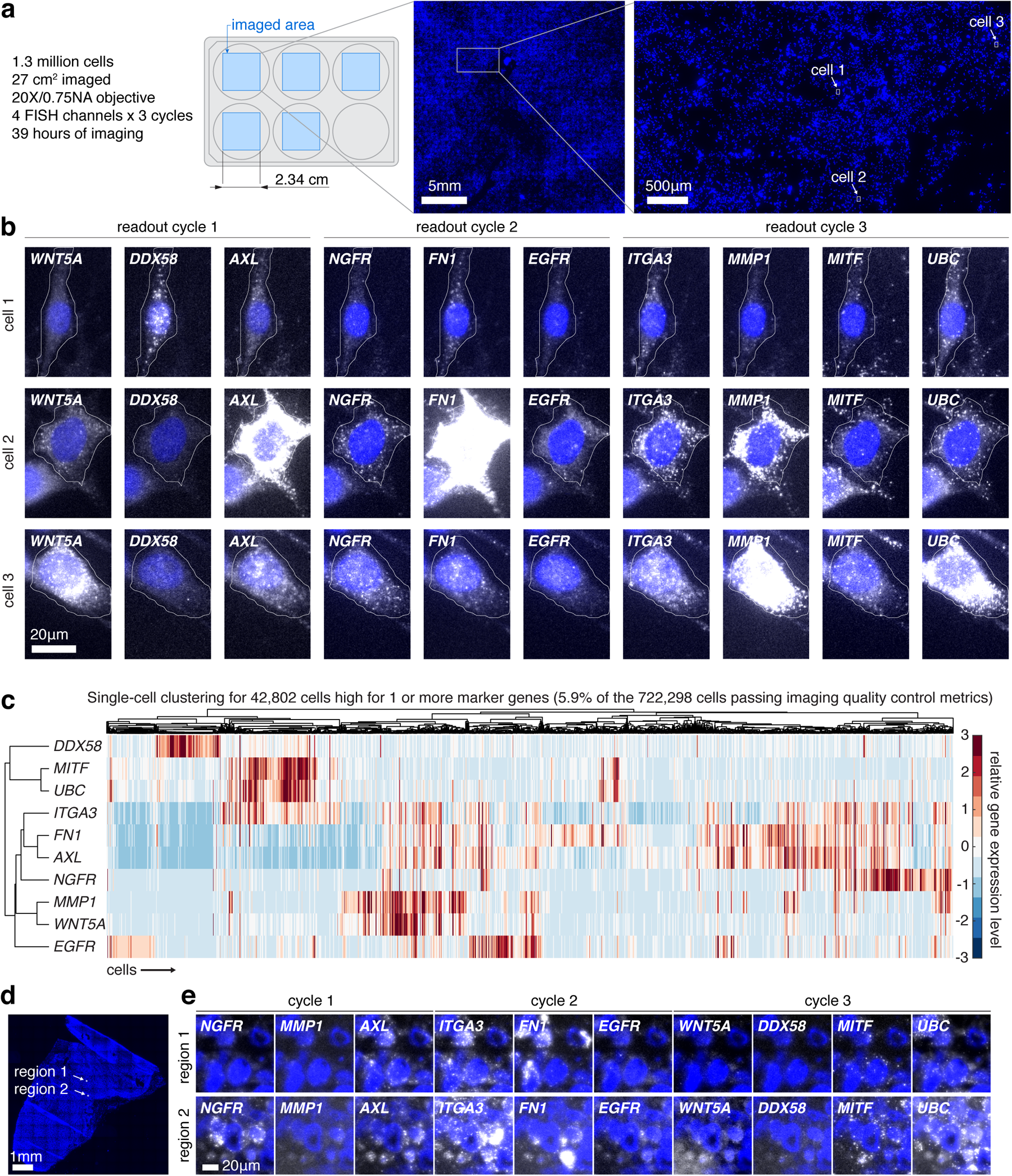
clampFISH 2.0 rapidly identifies rare cellular subpopulations in cell lines and tissue. **(a)** In the high-throughput profiling experiment, clampFISH 2.0 was performed for 10 genes in 1.3 million drug-naive WM989 A6-G3 cells, with **(b)** images at 20X magnification of 3 example cells with 10 genes probed throughout 3 readout cycles. **(c)** We detected 42,802 cells (5.9% of the 722,298 cells passing quality control checks) that expressed high levels of one or more of 8 cancer marker genes (*WNT5A*, *DDX58*, *AXL*, *NGFR*, *FN1*, *EGFR*, *ITGA3*, *MMP1*) and performed hierarchical clustering on this population. **(d)** A 20X magnification scan of DAPI in a fresh frozen tumor model with human WM989-A6-G3-Cas9-5a3 cells injected into a mouse that was fed chow containing a BRAF^V600E^ inhibitor. **(e)** 20X magnification images of clampFISH 2.0 spots in the same tissue section as in (d), probing for the same 10 genes as in (b).

As an additional measure of clampFISH 2.0’s quantitative performance, we compared the average clampFISH 2.0 spot count for ten genes with their relative abundance (transcripts per million) as detected by bulk RNA sequencing, and found moderate correlations in two melanoma cell lines (R^2^ between 0.256 and 0.607; see Supplementary Figure 9). We observed that *FN1* and *MMP1*, both of which have a lower mean clampFISH 2.0 spot count than would be expected from the remaining genes’ trend, are expressed at particularly high levels in a subset of cells (see Fig. 3b), suggesting that optical crowding at 20X magnification may contribute to their under-counting by clampFISH 2.0.

### Iterative hybridization enables multiplexed gene expression profiling of over one million cells

A crucial advantage of clampFISH 2.0 is its potential for rapid multiplexing through iterative hybridization of readout probes. Iterative hybridization refers to schemes for multiplexing beyond the spectral capabilities of conventional fluorescence microscopes (Lubeck et al. 2014). The basic idea is to detect RNA FISH signal from a small number (typically 3-4) of RNA targets using spectrally distinct fluorophores for each target. To measure RNA FISH signal from more targets in the same cells, the signal from the current set of targets is removed and then another round of hybridization to the next set of targets is performed, enabling detection of another set of RNA species. ClampFISH 2.0 is, in principle, ideally suited for such iterative schemes because all the scaffolds can be generated at once before any readout steps, and the short readout probes could be stripped and reprobed very rapidly.

An important first step for iterative hybridization is the ability to remove the fluorescent signal from the sample after imaging. Thus, we first tested whether the readout probes could be reliably stripped from their scaffolds with a simple high-stringency wash. We probed the mRNA from 10 genes, each with its own primary probe set with one of ten amplifier-specific sequences (pairing gene 1 with amplifier set 1, gene 2 with amplifier set 2, and so on), and generated scaffolds by amplifying to round 8. With these scaffolds generated in three separate wells, we then hybridized of 4 spectrally distinct sets of readout probes (coupled to Atto488, Cy3, Alexa Fluor 594, or Atto 647N), each binding to a specific amplifier set, thus visualizing four genes simultaneously per well (10 genes total, where scaffolds for one housekeeping gene, *UBC*, were probed in all 3 wells). After imaging these spots, we then stripped off the readout probes with 30% formamide in 2X SSC, re-imaged the samples, and noticed nearly all spots were removed (Supplementary Figures 10 and 11). Since the clampFISH 2.0 scaffolds are constructed of interlocking loops, we expected the scaffolds to remain stably attached to the mRNA targets despite the dissociation of the readout probes. Indeed, when we re-probed the same scaffolds after multiple cycles of readout hybridization and stripping, we observed the same spots as in the initial readout cycle (Supplementary Figures 12) and found high correlations in spot counts when compared to the initial readout cycle (eg. R^2^ of 0.833 (replicate 1) and 0.846 (replicate 2) for *WNT5A*; see Supplementary Figures 13 and 14), thus demonstrating the stability of the scaffolds and their ability to be repeatedly probed. In fact, even after leaving a sample refrigerated for 4 months and again re-probing the same scaffolds, we still observed the same spots (Supplementary Figure 12) and maintained a similarly high correlation in spot counts to the initial readout cycle (R^2^ of 0.758 (replicate 1) for *WNT5A; see* Supplementary Figure 13), thus demonstrating flexibility in the timing of readout and imaging.

Having demonstrated the ability to strip off readout probes, we then attempted to detect the mRNA from 10 different genes simultaneously in individual cells. We targeted the transcripts of the genes *WNT5A*, *DDX58*, *AXL*, *NGFR*, *FN1*, *EGFR*, *ITGA3*, *MMP1*, *MITF*, and *UBC* at the same time in the WM989 A6-G3 melanoma cell line (Shaffer et al. 2017) (and WM989 A6-G3 RC4 cells; see methods for details). We imaged cells spread over 5 wells of a 6-well culture dish with 3 cycles of imaging. Each imaging cycle consisted of detection in 4 readout probe channels, with *UBC* probed in every cycle as a control for consistency. The amplified signal allowed for a typical exposure time of 250ms with a 20X/0.75NA objective lens, allowing us to detect 10 genes in 1.3 million cells in 39 hours of imaging (Fig. 3a,b), demonstrating the ability to perform multiplex gene expression analysis via iterative hybridization across a large number of individual cells.

As a demonstration of the sorts of analyses that such high-throughput multiplexed RNA quantification enables, we analyzed the co-expression of these genes in the rare subpopulations that express them. Previous work has demonstrated that these genes are only expressed highly in rare cells (1:50-1:500), and that that it is these rare cells with high expression that are the ones that survive targeted drug therapies (Shaffer et al. 2017; Emert et al. 2021; Schuh et al. 2020). Many of these genes co-express in single cells (Shaffer et al. 2017), but the precise coexpression relationships have been hard to decipher due to the rarity of the expression. We reasoned that the much higher number of cells that we could image with multiplex clampFISH 2.0 (~1.3M vs. ~8700 for conventional single-molecule RNA FISH (Shaffer et al. 2017)) would enable us to measure these relationships. Using automated cell segmentation (Stringer et al. 2021) and a spot-detection pipeline, we identified 42,802 cells with one or more marker genes positively-associated with drug resistance out of a total pool of 722,298 cells. This sample size was large enough that we could observe distinct clusters of co-expression (Fig. 3c; for a technical replicate, see Supplementary Figure 15), including subsets of cells associated with distinct resistance phenotypes (Emert et al. 2021), demonstrating the sorts of analyses that are now possible with the high throughput of clampFISH 2.0.

### ClampFISH 2.0 detects RNA in tissue sections

An important application of image-based gene expression detection methods is in multicellular organisms and tissues. To demonstrate that clampFISH 2.0 could work in this context as well, we used the same 10-gene panel described above in fresh frozen tumor sections with 6µm thickness. These sections came from the injection of WM989-A6-G3-Cas9-5a3 cells into mice, which subsequently grew into tumors and were then treated with the BRAF^V600E^ inhibitor PLX4720 (samples first used in (Torre et al. 2021); see that paper for details). We observed clampFISH 2.0 signal in many of the cells, including consistent *UBC* signal in virtually all cells we assessed as of presumptive human origin but minimal signal in those of presumptive mouse origin (species distinguished by two distinct nuclear morphologies; see Supplementary Figure 17), confirming that clampFISH 2.0 was able to detect RNA in tissue sections (Fig. 3d,e). We also performed clampFISH 2.0 in a formalin-fixed paraffin embedded (FFPE) tissue section, in which we saw dimmer *UBC* clampFISH 2.0 signal (Supplementary Figure 16). There were regions of the FFPE tissue section that were completely devoid of signal, perhaps due to sample degradation or other unknown factors.

## Discussion

We have here described the development of an improved version of clampFISH, termed clampFISH 2.0. Its key features are the inverted probe design, which makes probe synthesis far more cost and time efficient, and the increased speed of the protocol. In particular, the efficiencies for probe synthesis are critical for multiplexing applications in which one targets multiple RNA species at the same time.

One important aspect of amplified signal is that one can use lower-powered optics, in particular at lower magnification. By using a 20X (or 10X) objective, we can obtain a 20-25 fold (40-75 fold) increase in throughput (number of cells imaged per unit time) as compared to conventional single-molecule RNA FISH imaged using a 60x objective. These order-of-magnitude increases in throughput can enable many new applications, especially in the detection of rare cell types. It is possible that other imaging improvements may be enabled by the dramatically increased signal afforded by signal amplification.

While here we have demonstrated a straightforward iterative hybridization scheme for multiplex RNA detection, one could imagine using clampFISH 2.0 for more complex combinatorial multiplex schemes as well (Lubeck et al. 2014; Shah, Lubeck, Schwarzkopf, et al. 2016; Shah, Lubeck, Zhou, et al. 2016; Eng et al. 2019; Moffitt, Hao, Wang, et al. 2016; Moffitt, Hao, Bambah-Mukku, et al. 2016; Xia et al. 2019). Many of those schemes rely on the detection of the same RNA in a specified subset of iterative detection rounds. ClampFISH 2.0 could be particularly well-suited for such schemes, because one could use combinations of readout probes in each round to detect specific RNA species. That readout probes can be re-hybridized to the same scaffolds offers added flexibility in sequential encoding schemes. For example, whereas the sequential barcode is normally encoded by the library of RNA-binding probes, which cannot be modified after their construction, each gene might instead have a single associated amplifier set, where the choice of each imaging cycle’s subset of readout probes would define the barcode.

Another potential benefit of clampFISH 2.0 for such sequential barcoding schemes is the small optical size of the spots, which generally appear to be at or near the diffraction limit. Both hybridization chain reaction and rolling-circle amplification produce spots that are larger (up to ~1µm) (Xia et al. 2019; Shah, Lubeck, Schwarzkopf, et al. 2016; J. H. Lee et al. 2015) than diffraction-limited spots, which contributes to optical crowding—if visualizing a large number of spots, they can overlap, making it difficult to discriminate neighboring spots. That makes it particularly difficult to colocalize spots through multiple rounds of hybridization and imaging. Other benefits of a diffraction-limited spot size is that the small size is beneficial for accurate super-resolution structural analysis by e.g. STORM (Rust, Bates, and Zhuang 2006), DNA-PAINT (Auer et al. 2017; J. Lee, Park, and Hohng 2018; Giannone et al. 2010; Sharonov and Hochstrasser 2006; Schoen et al. 2011), or STED (Hell 2003), and also that many image analysis tools assume diffraction-limited spots.

## Methods

### Probe design and construction

#### ClampFISH 2.0 primary probe design and construction

We constructed clampFISH 2.0 primary probes as follows. First, we designed a set of 30mer RNA-targeting probe sequences for each target gene with custom MATLAB software (Raj et al. 2008) and added a flanking 10mer 5’ sequence (AAGTGACTGT) and a 10mer 3’ sequence (ACATCATAGT) to each of those respective ends, producing a 50mer sequence (Supplementary Table 1). The 50mer sequences were run through a custom MATLAB script using BLAST (Camacho et al. 2009) for alignment to the human transcriptome and NUPACK (Dirks and Pierce 2004a; Dirks et al. 2007; Dirks and Pierce 2003; Fornace, Porubsky, and Pierce 2020) to predict binding energies of the off-target transcriptomic hits. We kept only the hits with binding energy less than −14 kcal/mol, and then assigned each of these hits the maximum fragments per kilobase of transcript per million (FPKM) from a set of 13 human RNA-seq datasets from the ENCODE portal (Davis et al. 2018; ENCODE Project Consortium 2012) (https://www.encodeproject.org/), see Supplementary Table 6 for file identifiers. For each gene, we selected 24-32 primary probes per gene target, with a preference for probes targeting the coding region and where the sum of FPKM values from its predicted off-target hits was minimized. For probes targeting GFP, we used 10 probes whose 30mer primary probe sequences were taken from (Rouhanifard et al. 2018). The 50mer sequences were ordered from Integrated DNA Technologies (IDT) and pooled together for a given gene. For each gene-specific pool, an azido-dATP (N6-(6-Azido)hexyl-3’-dATP, Jena Bioscience, NU-1707L) was added to the probes’ 3’ ends with Terminal Transferase (New England Biolabs, M0315L), which adds a single azido-dATP molecule. Then, the 5’ ends were phosphorylated with T4 Polynucleotide Kinase (New England Biolabs, M0201L). Each gene-specific pool of 51mer oligonucleotides was mixed with a 20mer ligation adapter (ACAGTCACTTCAACACTCAG) and a 58mer oligonucleotide, which were both ordered from IDT. The 58mer oligonucleotide was ordered with a 5’ alkyne modification (5’ hexynyl) and was designed with the following sequences, in 5’ to 3’ order: a universal 18mer sequence (AGACATTCTCGTCAAGAT), an amplifier-specific 30mer sequence (serving as a landing pad upon which a secondary probe can bind), and a universal 10mer sequence (CTGAGTGTTG). Then, T7 DNA Ligase (New England Biolabs, M0318L) was added, ligating together a complete 109mer (50 + 1 + 58) primary probe. We then added ammonium acetate to a 2.5M concentration, centrifuged twice at 17,000g where each time we pipetted all but the bottom 20μL of solution to a new tube, ethanol precipitated the probes, resuspended the probes in nuclease-free water, centrifuged the tube at 17,000g, and pipetted all but the bottom 5μL into a new tube. See Supplementary Tables 1, 2, 4, and 7 for sequences and information related to primary probes.

#### ClampFISH 2.0 amplifier probes design and construction

clampFISH 2.0 amplifier probes (secondary probes and tertiary probes) were constructed as follows. To design amplifier probe sets 1 and 2, two 30mer ‘landing pad’ sequences (one for the secondary, one for the tertiary) were manually generated with approximately 50% GC content and “AT” at the center, and the 30mer was then concatenated to itself to form a 60mer backbone sequence. We added 15mer arms on each end of the 60mer secondary backbones, such that arms were reverse complements to their paired tertiary backbone, and similarly added 15mer arms to each tertiary backbone to be reverse complements to their paired secondary backbone, thus completing each amplifier probe’s full 90mer probe sequence. For the remaining amplifier series, we generated 500,000 random 30mers, then replaced the middle two bases with “AT”. We kept sequences where the percent GC content of the left 15 nucleotides and the right 15 nucleotides were both between 45% and 55%, and then concatenated the remaining two 30mers together to create a 60mer backbone sequence. Backbone sequences with stretches of 3 or more C, 3 or more G, or 5 or more G or C bases were discarded. For amplifier series 3 to 7, we then selected backbones where the free energy of each backbone’s folded structure was greater than −2kcal/mol as predicted using the DINAMelt web server (Markham and Zuker 2005), selected those without hits against the human transcriptome using BLAST (https://blast.ncbi.nlm.nih.gov), added two 15mer arms to each backbone as before to generate a 90mer amplifier probes, and then selected the five 90mer amplifier probe pairs where the free energy of folding was the least negative as predicted using DINAMelt. For amplifier series 8 to 15, we followed the same steps to generate 60mer backbones (using a different random number generator seed), and then used NUPACK to predict the minimum free energy of its folded structure, accepting those with a value greater than −1.5 kcal/mol. We designated half the 60mer sequences to be secondary backbones and the other half to be tertiary backbones and paired each secondary backbone with a tertiary backbone. We again used NUPACK to keep only those with a minimum free energy greater than −2.0 kcal/mol. We checked for off-target binding against the human transcriptome using BLAST, both using a spliced transcriptome database and a custom-generated transcriptome database with unspliced transcripts, used NUPACK to keep only those with strong off-target binding to RNAs, then took the sum of the RNA transcripts’ maximum FPKM from the ENCODE RNA-seq datasets to generate an off-target FPKM for each secondary and tertiary probe. We chose secondary and tertiary probe pairs where each probes’ FPKM sum is ≤500 when using the spliced transcript database and ≤2500 when using the unspliced transcript database. We then dropped any amplifier sets with probes hitting genomic repeats using repeatmasker (https://www.repeatmasker.org/). We used NUPACK to simulate binding against other probes of the same probe type (each secondary against other secondaries, each tertiary against other tertiaries), and discarded 4 amplifier sets where the predicted binding energy to another probe was <-23 kcal/mol. All amplifier probe sequences are listed in Supplementary Table 2.

We ordered amplifier probes from IDT as 89mers with a 5’ hexynyl modification for 15 amplifier sets in total (15 secondaries and 15 tertiaries). In separate reactions for each amplifier probe, we added an azido-dATP (N6-(6-Azido)hexyl-3’-dATP, Jena Bioscience, NU-1707L) to the probes’ 3’ ends with Terminal Transferase, thus completing the 90mer amplifier sequence. We then added ammonium acetate to 2.5M and magnesium chloride to 10mM, then centrifuged twice at 17,000g where each time we pipetted all but the bottom 10μL of solution to a new tube. We then ethanol precipitated the probes, resuspended the probes in 200μL nuclease-free water, centrifuged the tube at 17,000g and pipetted all but the bottom 20μL into a new tube.

#### ClampFISH 2.0 readout probe design and construction

For the amplifier screen experiment, we used a 20 nucleotide readout probe that was designed to bind to the center of the 30mer landing pad sequences of each secondary probe. The readout probes were ordered from IDT with a 3’ Amino modifier (/3AmMO/), coupled to Atto 647N NHS-ester (ATTO-TEC, AD 647N-31), ethanol precipitated, purified by high-performance liquid chromatography (HPLC) (Raj et al. 2008), and resuspended in TE pH 8.0 buffer (Invitrogen, AM9849).

For all other experiments we designed two readout probes for each amplifier set: one to bind to the secondary probe, and one to bind to the tertiary probe, where each was designed to bind roughly to the center of the probe’s 30mer landing pad sequences. Readout probe sequences were chosen such that the Gibbs free energy of binding to their target amplifier backbone (DNA:DNA binding) was −22 kcal/mol or −24 kcal/mol, as calculated by MATLAB’s oligoprop function (based on the parameters from (Sugimoto et al. 1996)), and then ordered from IDT with a 3’ Amino modifier. The two readout probes targeting a given amplifier set were pooled together and then coupled to one of four NHS-ester dyes (Atto 488, ATTO-TEC, AD 488-31; Cy3, Sigma-Aldrich, GEPA23001; Alexa Fluor 594, ThermoFisher, A20004; or Atto 647N, ATTO-TEC, AD 647N-31), ethanol precipitated, purified by HPLC, and resuspended in TE pH 8.0 buffer, except for readout probes coupled to Atto 488 which were not pooled until after the HPLC steps. All readout probe sequences can be found in Supplementary Table 3.

#### Conventional single-molecule RNA FISH probes

We designed conventional single-molecule RNA FISH probes for GFP, *AXL*, *EGFR*, and *DDX58* as previously described (Raj et al. 2008), but selected a subset of probes not overlapping with the clampFISH 2.0 primary probes for these genes. We coupled probes to the NHS-ester dyes Cy3 (for *AXL*, *EGFR*, and *DDX58* probe sets) and Alexa Fluor 555 (Invitrogen, A-20009; for the *GFP* probe set). All conventional single-molecule RNA FISH probe sequences are available in Supplementary Table 5.

All oligonucleotide sequences are available in Supplementary Tables 1-5. Scripts used to generate probe sequences are available at (https://www.dropbox.com/sh/q51kmcphoyi9yi3/AAB4g1a6ODDHaphsvbmBJAy-a?dl=0).

### Cell culture and tissue processing

The WM989 A6-G3 human melanoma cell line, first described in (Shaffer et al. 2017) was derived from WM989 cells (a gift from the lab of Dr. Meenhard Herlyn) that were twice isolated from a single cell and expanded. WM989 A6-G3 H2B-GFP cells were derived by transducing WM989 A6-G3 cells with 60µL Lenti_EFS (https://benchling.com/s/seq-6Jv3Rmebv1nIevxPfYQ6/edit), isolating a single cell, and expanding this clone (Clone A11). Both lines were cultured in Tu2% media (80% MCDB 153, 10% Leibovitz’s L-15, 2% FBS, 2.4mM CaCl_2_, 50 U/mL penicillin, and 50 μg/mL streptomycin). WM989 A6-G3 RC4 cells were derived by treating WM989 A6-G3 cells with 1µM vemurafenib in Tu2%, isolating a single drug-resistant colony, and culturing these cells in 1µM vemurafenib in Tu2% (Goyal et al. 2021) for several months. All cell lines were passaged with 0.05% trypsin-EDTA (Gibco, 25300120).

For the amplifier screen and pooled amplification experiment, WM989 A6-G3 H2B-GFP and WM989 A6-G3 RC4 cells were mixed together and plated on coverslips (VWR, 16004-098, 24×050mm, No. 1 coverglass) with 24-well silicone isolators (Grace Bio-Labs, 665108). For the readout probe stripping experiment, conventional single-molecule RNA FISH comparison experiment, and the amplification characterization experiment, we plated WM989 A6-G3 or WM989 A6-G3 RC4 cells into separate wells of an 8-well chambers (Lab-tek, 155411, No. 1 coverglass). For the high-throughput profiling experiment, we plated WM989 A6-G3 cells into 5 wells and WM989 A6-G3 RC4 cells into 1 well of a 6-well plate (Cellvis, P06-1.5H-N, No. 1.5 coverglass), and allowed them to grow out for 6 days (2-3 cell divisions for WM989 A6-G3 cells) before fixation.

We fixed cell lines at room temperature by rinsing cells once in 1xPBS (Invitrogen, AM9624), incubating for 10 minutes in 3.7% formaldehyde (Sigma-Aldrich, F1635-500ML) in 1xPBS, then rinsing twice in 1xPBS. Cells were permeabilized in 70% ethanol and placed at 4°C for at least 8 hours. Nuclease-free water (Invitrogen, 4387936) was used in all buffers used for fixation onwards, including permeabilization, probe synthesis, and all RNA FISH steps.

For the fresh frozen tissue experiment, a melanoma xenograft tumor was taken from experiments described in (Torre et al. 2021). Briefly, human WM989-A6-G3-Cas9-5a3 cells (without a genetic knockout), derived by isolating and expanding a single WM989 A6-G3 cell, were injected into 8-week-old NOD/SCID mice (Charles River Laboratories) and fed AIN-76A chow (mouse #8947) or AIN-76A chow containing 417mg/kg PLX4720 (mouse #8948). Once the tumor reached 1,500mm^3^ the mouse was euthanized, and the tumor tissue was dissected and placed in a cryomold with optimal cutting temperature compound (TissueTek, 4583), frozen in liquid nitrogen, and then stored at −80°C. Tumors were then sectioned on a cryostat to 6μm thickness, placed onto a microscope slide (Fisher Scientific, 6776214), fixed and permeabilized with the same protocol used for cell lines while in LockMailer slide jars (Fisher Scientific, 50-340-92), and then stored at 4°C.

For the formalin-fixed paraffin embedded (FFPE) tissue experiment, a piece of about 3×3×3mm^3^ patient-derived tissue from either WM4505-1 cells or WM4298-2 cells were implanted into an NSG mouse. When the tumor was palpable, mice were continuously fed a diet with BRAF/MEK inhibitors (PLX4720 200ppm + PD-0325901 7ppm, chemical additive diet, Research Diets, New Brunswick, NJ). Tumor size was assessed once weekly by caliper measurements (length x width^2^/2). When the tumors reached 1,000mm^3^ or when necessary for animal welfare, the tumor was harvested and immediately placed in 10% neutral buffered formalin overnight (less than 48hrs), washed once with 1xPBS, and stored in 70% ethanol at room temperature.

Following the Wistar Institute Histotechnology facility’s standard protocol, the fixed tumor samples were embedded in paraffin, sectioned to 5μm thickness, and placed on a microscope slide. To avoid exposure to the air, the samples were sealed with a thin layer of paraffin, then stored at room temperature. Within 24 hours of beginning the clampFISH 2.0 steps, we performed a protocol adapted from <u>(Choi et al. 2018; Acheampong et al. 2022)</u>. First, we deparaffinized the sample in HistoChoice Clearing Agent (Sigma-Aldrich, H2779-1L) for 40 to 110 minutes total, then, while the sample was in LockMailer jars, in 100% ethanol for 2 x 5 minutes. We then post-fixed the sample in 3:1 methanol-acetic acid (v/v) solution for 5 minutes, re-hydrated the sample in nuclease-free water for 3 minutes, then performed an antigen retrieval step by placing the sample for 15 minutes into a LockMailer jar containing 10 mM sodium citrate pH6 with 0.1% diethyl pyrocarbonate (DEPC), which was heated in a ~100℃ water bath. We then stored the sample in 2X SSC for about 20 minutes at room temperature or overnight at 4℃.

For both the fresh frozen tissue and the FFPE tissue samples, we next placed the samples’ slides in 2X SSC for 1 - 5 minutes, in 8% sodium dodecyl sulfate (Sigma-Aldrich, 75746-250G; dissolved in nuclease-free water) for 2 minutes, and then into 2X SSC for up to 2 hours, after which we began the primary probe steps.

### ClampFISH 2.0 protocol

#### ClampFISH 2.0 primary probe steps

We performed clampFISH 2.0 in 8-well chambers as follows. First, we aspirated the 70% ethanol (or 2X SSC for tissue sections), rinsed with 10% wash buffer (10% formamide, 2X SSC), then washed with 40% wash buffer (40% formamide, 2X SSC) for 5-10 minutes. We mixed primary probes with 40% hybridization buffer (40% formamide, 10% dextran sulfate, 2X SSC) such that each probe’s final concentration was 0.1ng/μl (~2.8nM), added this mixture to the well, covered and spread it out with a coverslip, and then incubated overnight (10 or more hours) in a humidified container at 37°C. We hybridized only a single primary probe set per well with the amplifier screen experiment (GFP or *EGFR* probe sets) and the pooled amplification experiment (GFP probe set). For all other experiments we hybridized 10 primary probe sets together (see Supplementary Table 1 for list of primary probe sets).

The following day, we pre-warmed all wash buffers to 37°C: 10% wash buffer, 30% wash buffer (30% formamide, 2X SSC), and 40% wash buffer. We first added warm 10% wash buffer, removed the coverslips and aspirated the solution, and washed again with warm 10% wash buffer. We then washed 2 x 20 minutes with warm 40% wash buffer on a hotplate set to 37°C (the temperature setting used throughout the protocol). After removing the chamber from the hotplate, we added 10% wash buffer before beginning the amplification steps.

#### ClampFISH 2.0 amplification steps

For amplification, we first mixed together all the secondary probes with 10% hybridization buffer with Triton-X (10% formamide, 10% dextran sulfate, 2X SSC, and 0.1% Triton-X (Sigma-Aldrich, T8787-100ML)) to a final ~20nM concentration per probe (range: ~13nM to 25nM) with a 27mer circularizer oligonucleotide (TCTTGACGAGAATGTCTTACTATGATG) at a 40nM final concentration. We also mixed together all tertiary probes with 10% hybridization buffer with Triton-X at the same concentrations, but without the circularizer oligonucleotide. In preparation for multiple click reaction steps, we prepared tubes each with an appropriate volume of pre-warmed 2X SSC with Triton-X and DMSO (2X SSC, 0.25% Triton-X, 10% dimethyl sulfoxide) for the amplification step, and warmed them to 37°C. We also aliquoted sodium ascorbate (Acros, AC352680050) into 1.5mL tubes, ready to be dissolved fresh with each click step. We prepared a CuSO_4_ (Fisher Scientific, S25289) and BTTAA (Jena Bioscience, CLK-067-100) mixture in a 1:2 CuSO_4_:BTTAA molar ratio, enough to use for all the click reactions throughout the rounds of amplification.

We added the secondary probe and circularizer oligo-containing 10% hybridization buffer with Triton-X to the well, covered it with a coverslip, and incubated for 30 minutes in a 37°C incubator. After taking the chamber out of the incubator, we added warm 10% wash buffer, removed the coverslips, washed 2 x 1 minute with warm 10% wash buffer, and then washed again with warm 10% wash buffer for 10 minutes on the hotplate. We then took the chamber off the hotplate and added 2X SSC before the click reaction. The click reaction mixture was then prepared by first mixing the CuSO_4_ and BTTAA mixture with the pre-warmed 2X SSC with

Triton-X and DMSO buffer. Working quickly, we added nuclease-free water to an ascorbic acid aliquot and vortexed it until dissolved. We aspirated the 2X SSC solution from the well plate and quickly added aqueous sodium ascorbate to the CuSO_4_ + BTTAA + 2X SSC with Triton-X and DMSO mixture (final concentrations: 150μM CuSO_4_, 300μM BTTAA, 5mM sodium ascorbate, ~2X SSC, ~0.25% Triton-X, ~10% DMSO) and briefly mixed it by swirling the tube by hand. We immediately added the click reaction solution to the wells and incubated it on the hotplate for 10 minutes. Next, we aspirated the click reaction mixture and washed the sample with warm 30% wash buffer for 5 minutes on the hotplate. The above steps (amplifier probe hybridization, 10% wash buffer steps, click reaction, and 30% wash buffer step) constitutes a single round of amplification, and takes about 1 hour when accounting for pipetting time.

Before beginning the next round of amplification, we then replaced the 30% wash buffer with warm 10% wash buffer. (If, alternatively, a breakpoint was needed in between rounds of amplification, we instead replaced the 30% wash buffer with 2X SSC and stored the sample at room temperature for up to 2 hours or at 4°C for up to a day). We then proceeded to the next round of amplification, this time using tertiary probes instead of secondary probes. We dub the completion of the primary step as having performed clampFISH 2.0 to “round 1”, the first secondary step as “round 2”, the first tertiary step as “round 3”, the next secondary step as “round 4”, and so on. We ran all amplifications to round 8, involving 1 primary probe round and 7 amplification rounds, except where noted differently. At the end of the last amplification round, we placed the sample at 4°C in 2X SSC until the readout probe steps (typically the samples were stored overnight for readout and imaging the subsequent day).

For the amplifier screen experiment and the pooled amplification experiment, we proceeded with conventional single-molecule RNA FISH per (Raj et al. 2008) by first rinsing briefly with 10% wash buffer, adding GFP or *EGFR* probes as well as a 20 nucleotide secondary-targeting readout probe at 4nM final concentration in 10% hybridization buffer (10% formamide, 10% dextran sulfate, 2X SSC), covering with a coverslip, placing in a humidified container and incubating overnight in at 37°C, adding 10% wash buffer to remove the coverslip, washing 2 x 30 minutes in 10% wash buffer in a 37°C incubator, while adding 50 ng/mL of the nuclear stain 4′,6-diamidino-2-phenylindole (DAPI) to the second wash, after which we did not carry out further readout probe steps. For the wash and click steps that use a hotplate, in these two experiments we instead used a 37°C incubator or bead bath, with the sample in a LockMailer slide jar submerged in the appropriate buffer.

For the high-throughput profiling experiment in a 6-well plate, we replaced the use of a hotplate with a 37°C incubator and increased the incubation time of the 10 minute wash in 10% wash buffer, the 10 minute click reaction, and all steps in 30% wash buffer by an additional 4 minutes to accommodate the longer time to warm up.

#### Readout cycle steps

The following day, either directly following amplification or the subsequent conventional RNA FISH, we performed a readout probe cycle as follows. First, samples were brought to room temperature and rinsed once with room-temperature 2X SSC. For each amplifier set (each of which corresponds to a particular gene target) to be probed, two readout probes were hybridized (with one binding the secondary and one binding to the tertiary), both coupled to the same fluorescent dye. A set of readout probes for each of four spectrally distinguishable dyes could be included in a given readout cycle. Each readout probe was hybridized at a 10nM final concentration in 5% ethylene carbonate hybridization buffer (5% ethylene carbonate, 10% dextran sulfate, 2X SSC, 0.1% Triton-X) for 20 minutes at room temperature. We then aspirated the solution, washed 2 x 1 minute with 2X SSC with Triton-X (2X SSC, 0.1% Triton-X), 1 minute with 2X SSC buffer, 5 minutes with 2X SSC with 50 ng/mL DAPI, then replaced with 2X SSC before imaging.

After imaging a given readout cycle, we stripped off the readout probes by incubating 2 x 5 minutes at 37°C with 30% wash buffer pre-warmed to 37°C, then added 2X SSC before starting another readout cycle. If we imaged the post-strip sample, we incubated 5 minutes with 2X SSC with 50 ng/mL DAPI, and replaced the solution with 2X SSC before imaging.

For the conventional single-molecule RNA FISH comparison experiment, after stripping the readout probes we proceeded with conventional single-molecule RNA FISH, as described above, but instead with probes for *AXL*, *EGFR*, or *DDX58* without any additional readout probes.

### Imaging and image analysis

#### Imaging

For imaging we used a Nikon Ti-E inverted microscope equipped with an ORCA-Flash4.0 V3 sCMOS camera (Hamamatsu, C13440-20CU), a SOLA SE U-nIR light engine (Lumencor), and a Nikon Perfect Focus System. We used 60X (1.4NA) Plan-Apo λ (Nikon, MRD01605), 20X (0.75NA) Plan-Apo λ (Nikon, MRD00205), and 10X (0.45NA) Plan-Apo λ (Nikon, MRD00105) objective and filter sets for DAPI, Atto 488, Cy3, Alexa Fluor 594, and Atto 647N (see Supplementary Table 8 for filter sets used; filter sets can also be viewed at https://www.fpbase.org/microscope/455WNQygW6268avMhrTNx8/). All 60X images were taken using 2×2 camera binning, while 20X and 10X images used 1×1 binning.

#### Image analysis

All data and scripts used are all publicly accessible in a Dropbox folder (https://www.dropbox.com/sh/q51kmcphoyi9yi3/AAB4g1a6ODDHaphsvbmBJAy-a?dl=0), which use functions from rajlabimagetools (https://github.com/arjunrajlaboratory/rajlabimagetools) and Dentist2 (https://github.com/arjunrajlaboratory/dentist2/tree/clamp2paper) repositories for spot processing and thresholding.

For the amplifier screen experiment, we segmented cells in rajlabimagetools, manually selected minimum spot intensity thresholds for conventional single-molecule RNA FISH, and counted the spots above this threshold from a 60X magnification z-stack for each cell. For cells in which this count was 20 or greater, we took an equivalent number of the highest-intensity clampFISH 2.0 spots from that cell and used this list of clampFISH 2.0 spot intensities for plotting in Supplementary Figure 5 and for calculation of the median intensity in Supplementary Figure 6.

For the pooled amplification experiment (Supplementary Figure 7), in order to quantify the typical spot intensity we used rajlabimagetools to extract the 10,000 highest-intensity GFP clampFISH 2.0 spots from 60X z-stacks of 40 segmented cells per condition (an average of 250 spots per cell). The highest-intensity spots were chosen to eliminate potential biases associated with manually-chosen thresholds.

For the readout probe stripping experiment (Supplementary Figure 8), we segmented 39-48 cells in the before-stripping 20X images, chose gene-specific clampFISH 2.0 spot intensity thresholds, aligned the same segmentations to the post-stripping images, and extracted spot counts from the post-stripping images.

For the amplification characterization experiment, we used Cellpose (Stringer et al. 2021) to automatically segment cells using cellular background fluorescence in the YFP channel (with the DAPI channel also included as a Cellpose input), and excluded abnormally small or large cells. For each of the 4 probed genes we used rajlabimagetools to extracted the top N spots from each round of amplification, where: N = (number of cells)×k, and k is the assumed average number of spots per cell (k = 120, 1, 20, and 80 spots/cell for *UBC*, *ITGA3*, *FN1*, and *MITF*, respectively). To avoid saturating the camera’s photon-collecting capacity at higher rounds of amplification, we extracted spots from longer exposure times on amplification rounds 1,2, and 4 (1000, 1000, 500, and 500 milliseconds for each gene, respectively) and shorter exposure times on amplification rounds 6, 8, and 10 (all were 100 milliseconds), and scaled these intensities by the ratio of median spot intensities between the two exposure times at round 6. For all no-click conditions, we used the longer exposure times to extract spot intensities. We then normalized the data by dividing all intensity values by the median value from round 1, using these in Figure 1e, Supplementary Figure 3, and Supplementary Figure 4).

For the conventional single-molecule RNA FISH comparison experiment (Fig. 2a,b), we manually segmented cells from 60X images using rajlabimagetools, manually selected minimum spot intensity thresholds for the conventional single-molecule RNA FISH data, and counted spots in each cell from 11 z-planes at 0.5μm spacing. We scaled and aligned these segmentations to the 20X and 10X images, and extracted clampFISH 2.0 spots exceeding a gene-specific threshold for 20X (3 z-planes at 1μm spacing) and 10X (3 z-planes at 2μm spacing) images. To quantify sensitivity and specificity on the lowly-expressed gene *DDX58*, we denoted cells with 3 or more spots as *‘DDX58* high’ and with 2 or fewer spots as ‘*DDX58* low’, and did so using conventional single-molecule RNA FISH at 60X magnification (the gold standard) and using clampFISH 2.0 at 20X magnification. In two biological replicates (different passages of WM989 A6-G3 cells), we found clampFISH 2.0 at 20X magnification could identify *‘DDX58* low’ cells with a specificity of 97% (32/33 cells, replicate 1) and 99% (86/87 cells, replicate 2) and *‘DDX58* high’ cells with a sensitivity of 41% (35/86 cells, replicate 1) and 53% (10/19 cells, replicate 2).

For the high-throughput profiling experiment, we stitched and registered the tiled scans from multiple imaging cycles at 20X magnification using custom scripts (publically available via https://www.dropbox.com/sh/q51kmcphoyi9yi3/AAB4g1a6ODDHaphsvbmBJAy-a?dl=0) and then divided the scan into smaller subregions. We imaged 5 wells (replicate 1) and 1 well (replicate 2) of WM989 A6-G3 cells, dividing those scans into 10×10 subregions, and 1 well (replicates 1 and 2) of WM989 A6-G3 RC4 cells, dividing those scans into 6×6 subregions. We used Dentist2 to choose spot intensity thresholds, extract spots, and then assign those spots to cellular segmentations generated by Cellpose based on cellular background fluorescence (eg. autofluorescence) in the YFP channel (using the diameter parameter of 90 pixels for WM989 A6-G3 cells and 350 pixels for WM989 A6-G3 RC4 cells). We used the housekeeping gene *UBC*, for which a readout probe was hybridized on every readout cycle, for the following quality control steps. First, we kept only subregions where there was an average of at least 25 *UBC* spots per cell for all readout cycles (we observed that near the edges of the wells, fewer spots above our chosen thresholds were detected, presumably because the coverslip used to spread out all probe-containing solutions were smaller than the full well). We then took only cells where, for all readout cycles, the *UBC* spot count was: at least 4, at least 0.025/um^2^×cell area, always within 50% of the median count from all readout cycles. Out of the initial 1,297,062 (replicate 1) and 253,662 (replicate 2) WM989 A6-G3 cells segmented, 722,298 (replicate 1) and 234,410 (replicate 2) cells passed all quality control metrics and were included in downstream analyses. To analyze only cells expressing high levels of one or more of 8 marker genes, we chose, for each gene, the following minimum spot count thresholds (format: minimum spot count to be considered high-expressing, percentage of cells high-expressing in replicate 1): *WNT5A* (>=15, 0.59%), *DDX58* (>=10, 0.56%), *AXL* (>=25, 3.56%), *NGFR* (>=30, 1.07%), *FN1* (>=100, 2.79%), *EGFR* (>=5, 1.40%), *ITGA3* (>=50, 2.31%), *MMP1* (>=40, 1.48%). For the 5.93% of cells (42,802 out of 722,298) in replicate 1 expressing high levels of one or more marker genes, we used MATLAB’s clustergram function to perform hierarchical clustering (Fig. 3c).

For Supplementary Figures 11–14, we ran the same pipeline on a smaller imaged area in Well A1 that, in addition to the three readout cycles included previously, also included a re-imaging of readout cycle 1 and readout cycles 4 and 5 (both of which re-used the same readout probes from readout cycle 1). To define spots, a single minimum spot intensity threshold was chosen for each gene on each round. Thresholds for readout cycle 4 images were made the same as those in cycle 1. For readout cycle 5 (performed after storing the sample at 4°C for 4 months), the thresholds were increased by 67% to 83% (the cycle 5 signal presumably appeared brighter due to changes in the microscope’s optical path, i.e. greater sample illumination or increased transmission to the sensor).

### RNA sequencing

We performed bulk RNA sequencing as described in (Goyal et al. 2021). We conducted standard bulk paired-end (37:8:8:38) RNA sequencing using RNeasy Micro (Qiagen, 74004) for RNA extraction, NEBNext Poly(A) mRNA Magnetic Isolation Module (NEB E7490L), NEBNext Ultra II RNA Library Prep Kit for Illumina (NEB, E7770L), NEBNext Multiplex Oligos for Illumina (Dual Index Primers Set 1) oligos (NEB, E7600S), and an Illumina NextSeq 550 75 cycle high-output kit (Illumina, 20024906), as previously described (Mellis et al. 2021; Shaffer et al. 2017). Prior to extraction and library preparation, the samples were randomized to avoid any experimental and human biases. We aligned RNA-seq reads to the human genome (hg19) with STAR v2.5.2a and counted uniquely mapping reads with HTSeq v0.6.1 (Dobin et al. 2013; Mellis et al. 2021; Shaffer et al. 2017) and outputs count matrix. The counts matrix was used to obtain tpm and other normalized values for each gene using scripts provided at: https://github.com/arjunrajlaboratory/RajLabSeqTools/tree/master/LocalComputerScripts

## Author Contributions

IPD designed, performed, and analyzed all experiments, supervised by AR. BE and CLJ assisted with FISH protocol development. YG derived and isolated the WM989 A6-G3 RC4 cell line, assisted with tissue sectioning, assisted with cell culture, and performed all RNA sequencing. BE derived and isolated the WM989 A6-G3 H2B-GFP cell line. AK, CLJ, BE, and SHR assisted with probe synthesis protocol development. JL performed cell segmentation for the amplifier screen experiment. GMA, MEF and ATW performed the mouse experiment with WM989-A6-G3-Cas9-5a3 cells and provided dissected tumor samples for the fresh frozen tissue experiment. MX and MH performed the mouse experiment with patient-derived tissue and prepared the samples up to paraffin embedding.

## Acknowledgements

We thank Allison Coté and Ian A. Mellis for helpful input on image analysis. We thank the ENCODE Consortium and the lab of J. Michael Cherry for RNA-seq datasets used for probe design. We thank the Penn Center for Musculoskeletal Disorders Histology Core (P30 AR069619) for their guidance on tissue cryo-sectioning. We are grateful to the Wistar Institute’s Histotechnology facility (supported by Cancer Center Support Grant P30 CA010815), for processing the FFPE tissue samples. BE acknowledges support from NIH F30 CA236129, NIH T32 GM007170, and NIH T32 HG000046; YG holds a Career Award at the Scientific Interface from BWF, and acknowledges additional support from the Schmidt Science Fellowship in partnership with the Rhodes Trust; CLJ acknowledges support from NIH training grants F30 HG010822, T32 DK007780, and T32 GM007170; AK acknowledges support from NIH K00-CA-212437-02; MEF and ATW acknowledge support from R01CA174746 and R01CA207935; ATW is also supported by a Team Science Award from the Melanoma Research Alliance and P01 CA114046; MH acknowledges support from NIH grants RO1 CA238237, U54 CA224070, PO1 CA114046, P50CA174523 and the Dr. Miriam and Sheldon G. Adelson Medical Research Foundation; AR acknowledges support from 5-U2C-CA-233285-04, NIH 4DN U01 HL129998, NIH 4DN U01DK127405, NIH Center for Photogenomics (RM1 HG007743), NIH Director’s Transformative Research Award R01 GM137425, and the Penn Epigenetics Institute.

## Disclosure of Potential Conflicts of Interest

AR receives patent royalties from LGC/Biosearch Technologies related to Stellaris mRNA FISH products. AR is on the scientific advisory board of Spatial Genomics. ATW is on the Board of Directors at ReGAIN therapeutics. The remaining authors state no conflict of interest.

## Supplementary Tables

See supplementary Tables 1-8.

---

**Supplementary Figure 1:**
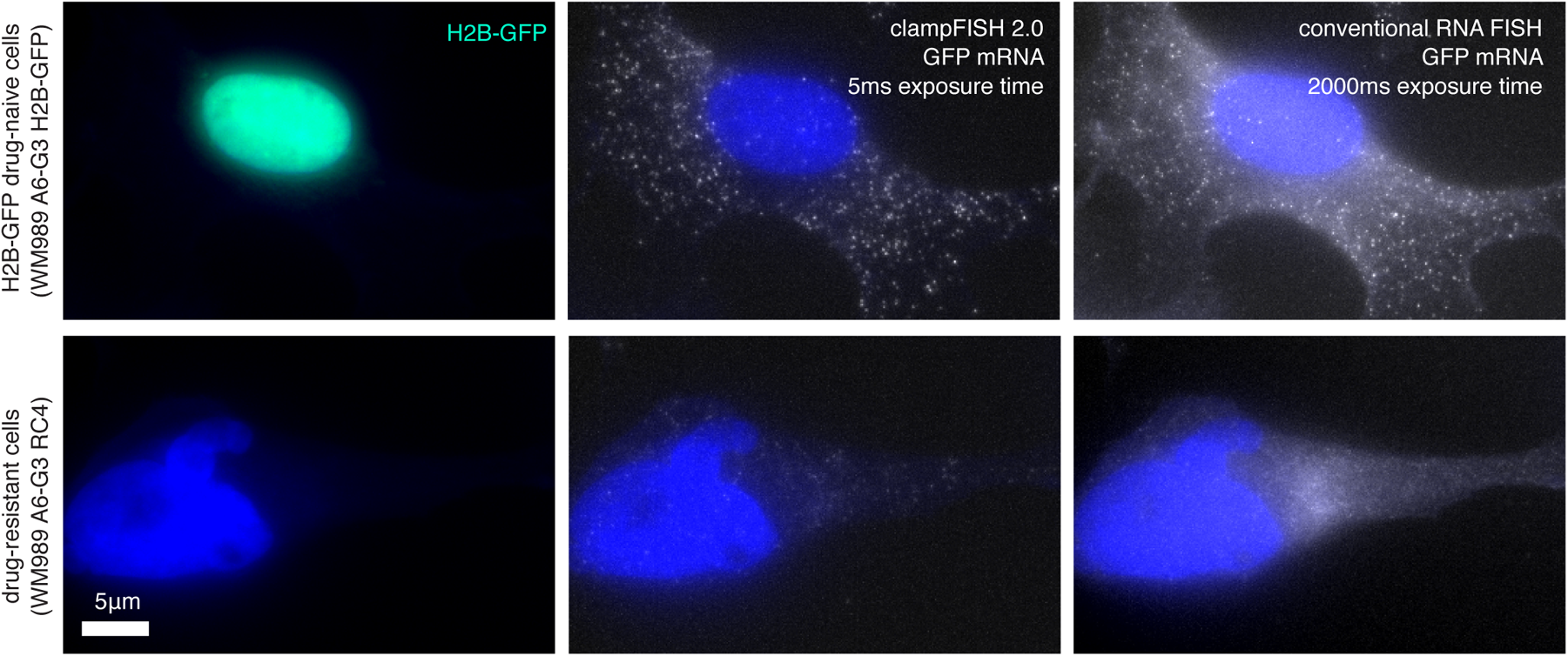
clampFISH 2.0 amplifies GFP RNA FISH signal. Images of GFP clampFISH 2.0 spots in drug-naive H2B-GFP WM989 A6-G3 cells (top) and vemurafenib-resistant WM989 A6-G3 RC4 cells (bottom) with a 20 nucleotide secondary-targeting readout probe (labeled with Atto 647N) and conventional single-molecule RNA FISH probes (labeled with Alexa 555) targeting different regions of the same RNA. Shown are maximum intensity projections of 9 z-planes at 60X magnification. As expected, we observed bright GFP clampFISH 2.0 spot counts in cells with nuclear-localized GFP signal, but not in cells without the H2B-GFP construct.

**Supplementary Figure 2:**
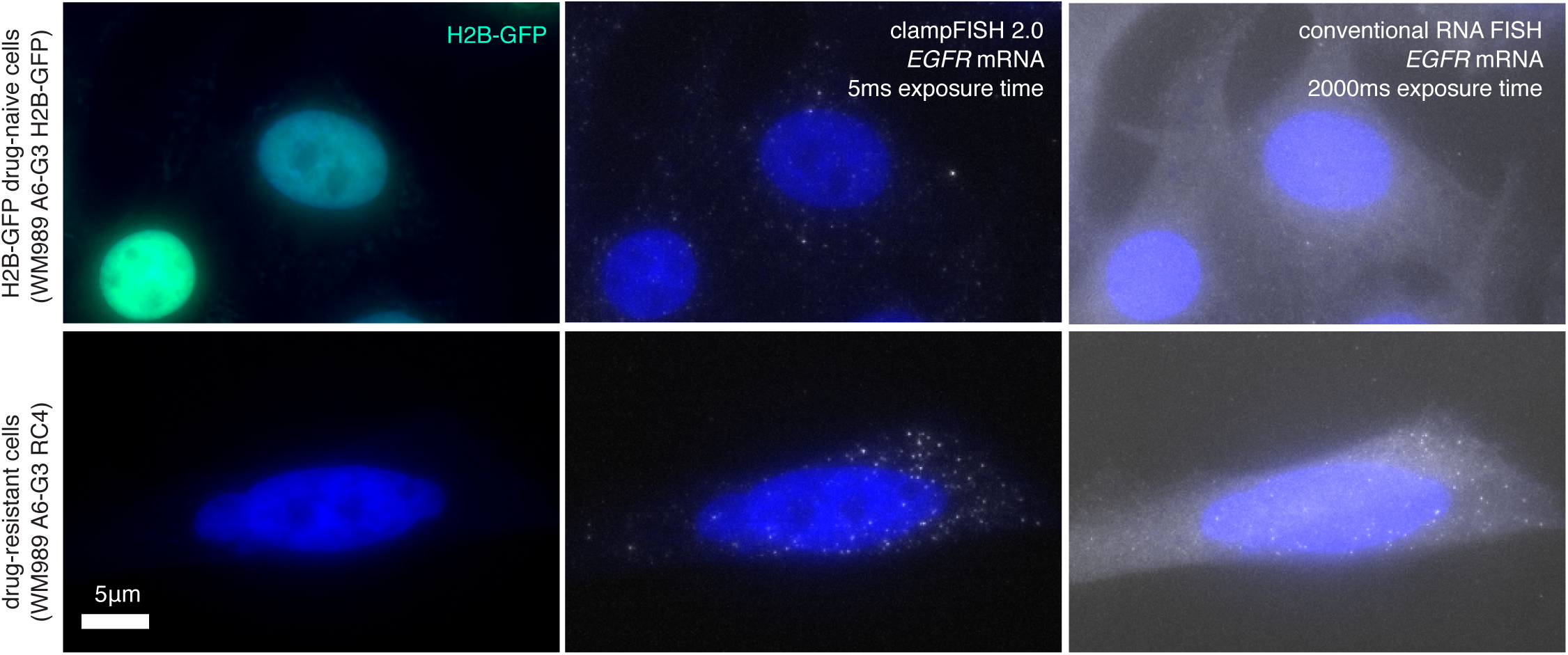
clampFISH 2.0 amplifies *EGFR* RNA FISH signal. Images of *EGFR* clampFISH 2.0 spots in drug-naive H2B-GFP WM989 A6-G3 cells (top) and vemurafenib-resistant WM989 A6-G3 RC4 cells (bottom) with a 20 nucleotide secondary-targeting readout probe (labeled with Atto 647N) and conventional single-molecule RNA FISH probes (labeled with Cy3) targeting different regions of the same RNA. Shown are maximum intensity projections of 9 z-planes at 60X magnification. As expected from bulk RNA-sequencing data, we observed many more *EGFR* clampFISH 2.0 spots in the vemurafenib-resistant cells than in the drug-naive cells.

**Supplementary Figure 3:**
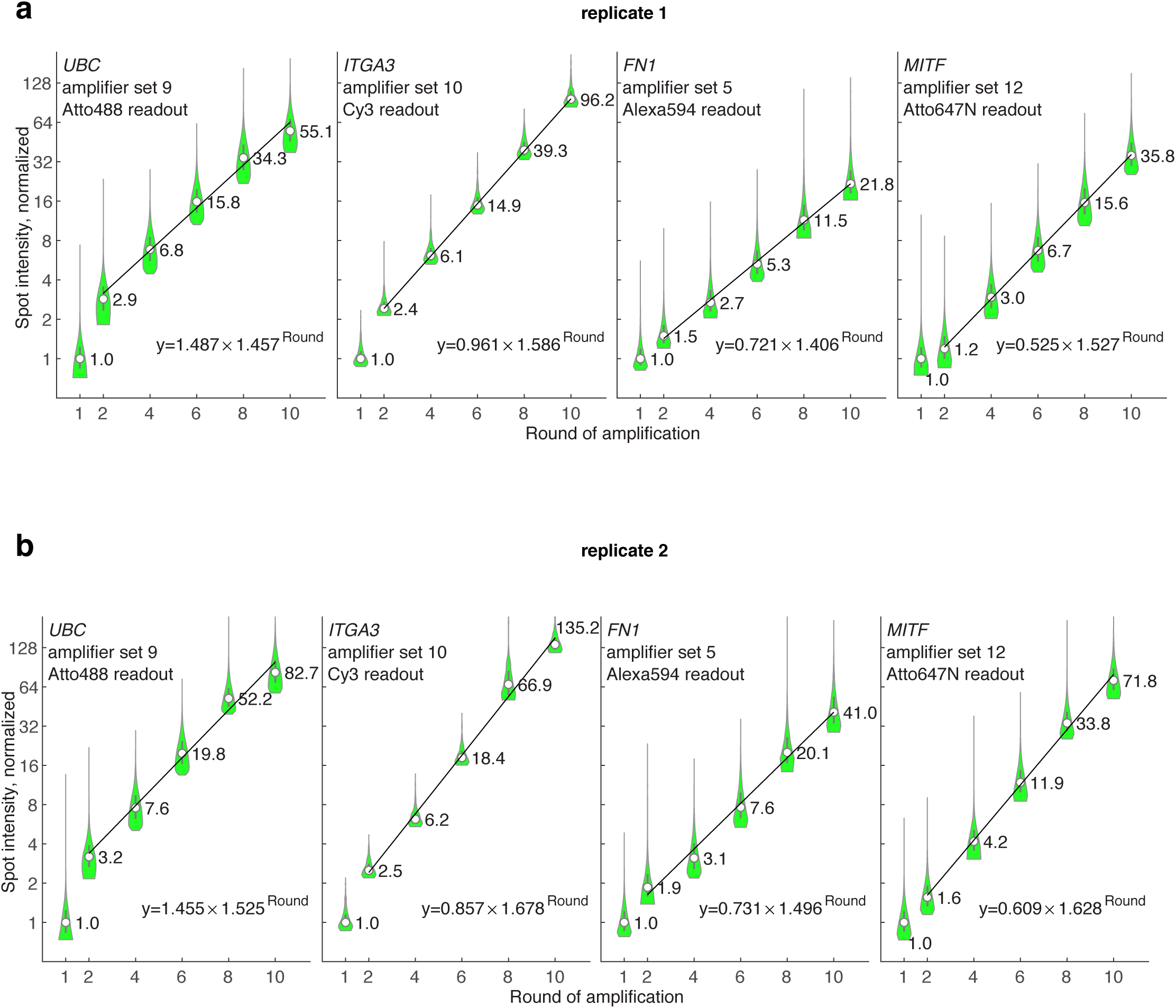
clampFISH 2.0 amplifies signal exponentially. **(a)** In an amplification characterization experiment, we performed clampFISH 2.0 with amplification to varying rounds (round 1, 2, 4, 6, 8, and 10) and then hybridized four readout probes to measure the spot intensities, with the median intensity from rounds 2, 4, 6, 8 and 10 fit to an exponential curve (labeled values are median intensities). We found that every round the spot intensities grew by a factor of 1.457, 1.586, 1.406, and 1.527 for each probe set respectively. With a hypothetical 2:1 binding ratio of each amplifier probe to the previous probe, these factors suggest a per-probe binding efficiency of 73%, 79%, 70%, and 76%, respectively. **(b)** Replicate 2 of the same experiment as in (a), where the spot intensities grew by a factor of 1.525, 1.678, 1.496, and 1.628, suggesting per-probe binding efficiencies of 76%, 84%, 75%, and 81%, respectively.

**Supplementary Figure 4:**
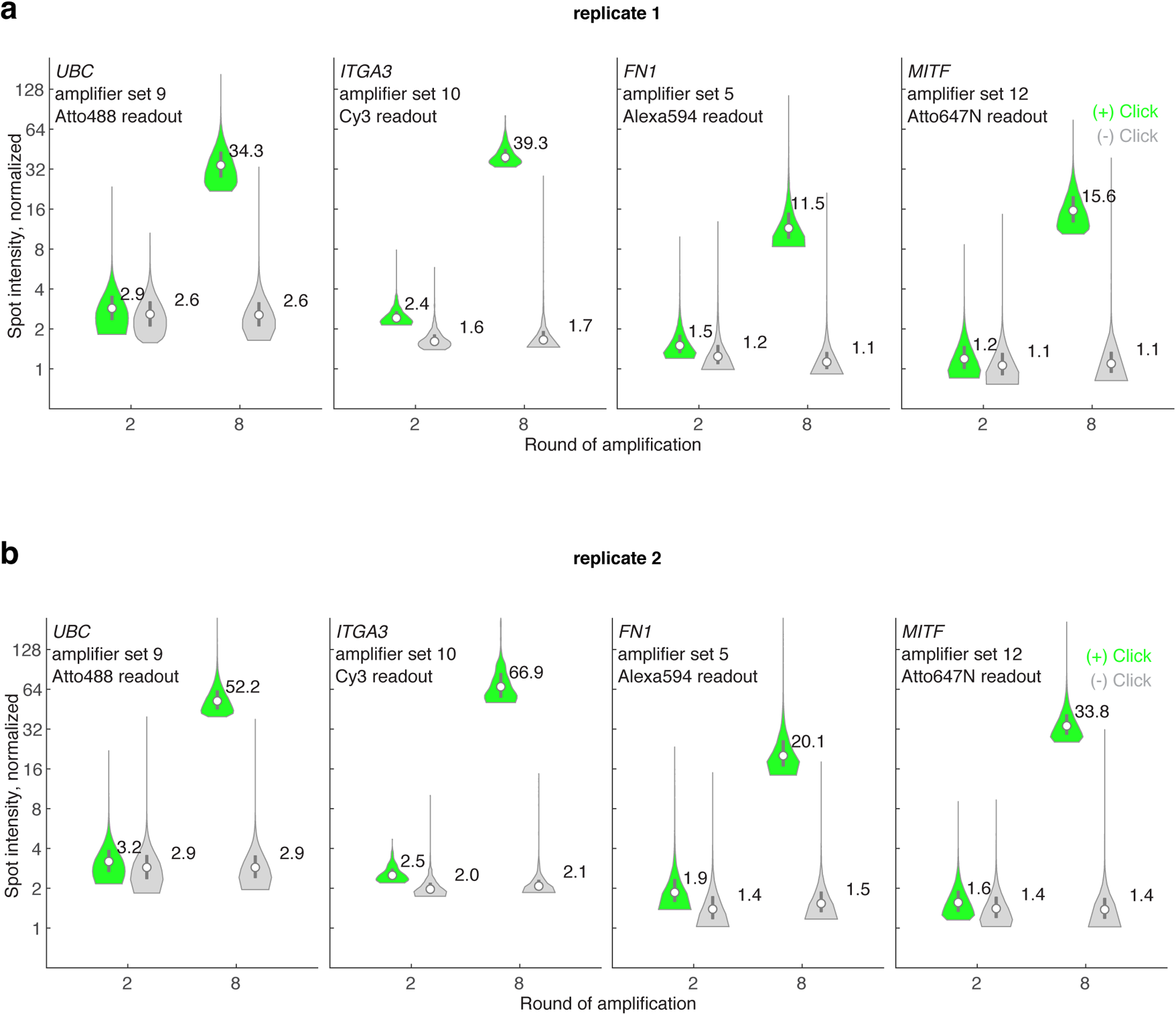
clampFISH 2.0 signal amplification is dependent on the click reaction. **(a)** In an amplification characterization experiment, we performed the clampFISH 2.0 amplification steps to rounds 2 and 8, both with (green) and without (grey) the copper sulfate catalyst included in the click reaction (labeled values are median intensities). **(b)** a biological replicate (different passage) of the same experiment as in (a). We saw no amplification from round 2 to round 8 in the absence of the copper catalyst, confirming that the click reaction is an essential step for clampFISH 2.0.

**Supplementary Figure 5:**
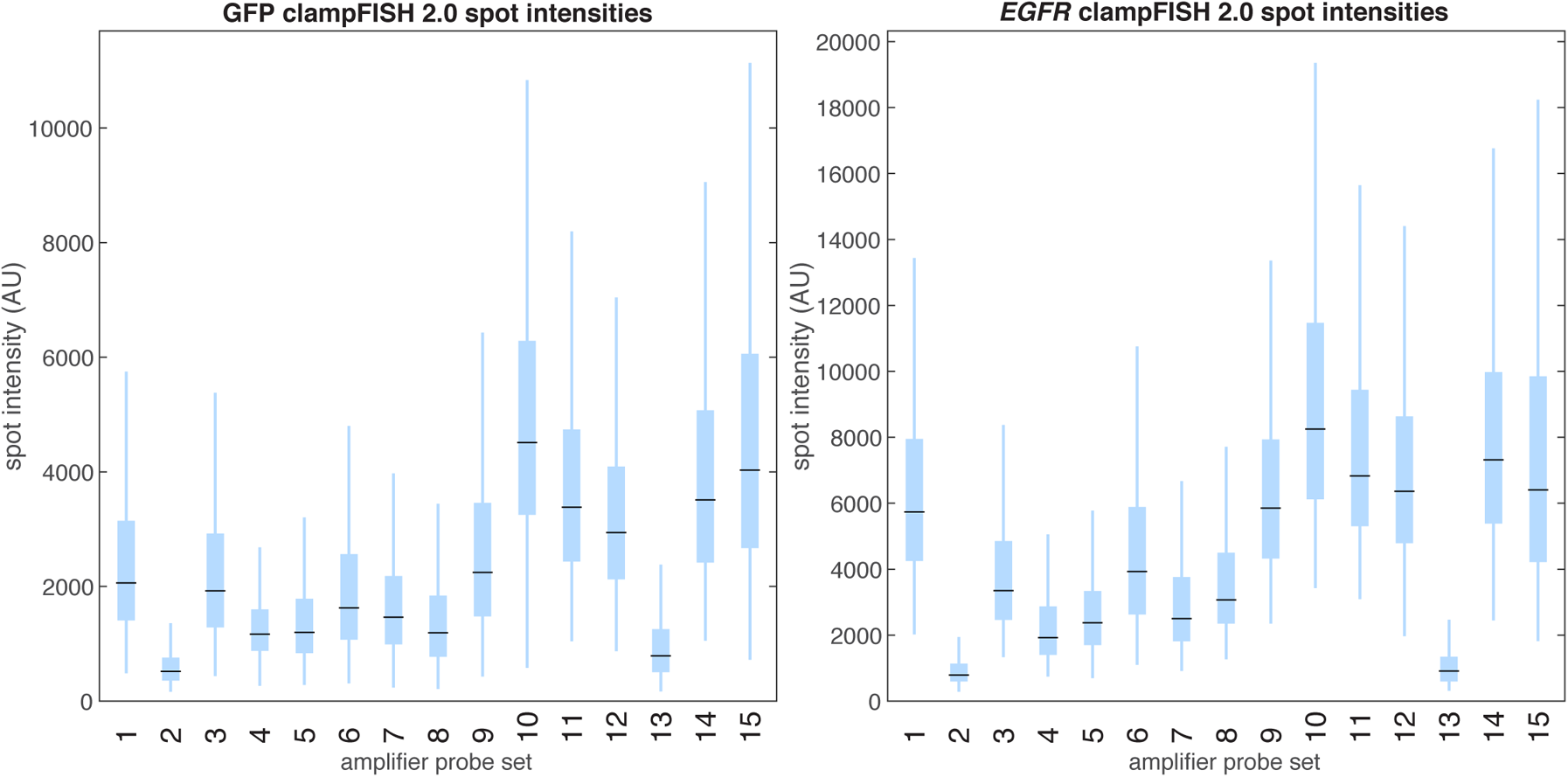
A screen of amplifier probe sets revealed designs with a high level of signal amplification. In a screen for amplifier probe sequences, we tested 15 amplifier sets, each used with primary probes targeting GFP (left) and *EGFR* (right) amplified to round 8 (in total, 30 primary probe sets were used: 2 targets x 15 amplifier sets). We labeled the clampFISH 2.0 scaffolds with 20 nucleotide secondary-targeting readout probes (coupled to Atto 647N) and performed conventional single-molecule RNA FISH (GFP probes in Alexa 555, *EGFR* probes in Cy3) targeting non-overlapping regions of the same mRNA as the primary probes. We counted the number of conventional single-molecule RNA FISH spots in each each segmented cell, took an equivalent number of the highest-intensity clampFISH 2.0 spots from that cell, and plotted these clampFISH 2.0 spot intensities (11,252 GFP and 881 *EGFR* outliers, out of 294,220 and 22,861 total points respectively, are not shown).

**Supplementary Figure 6:**
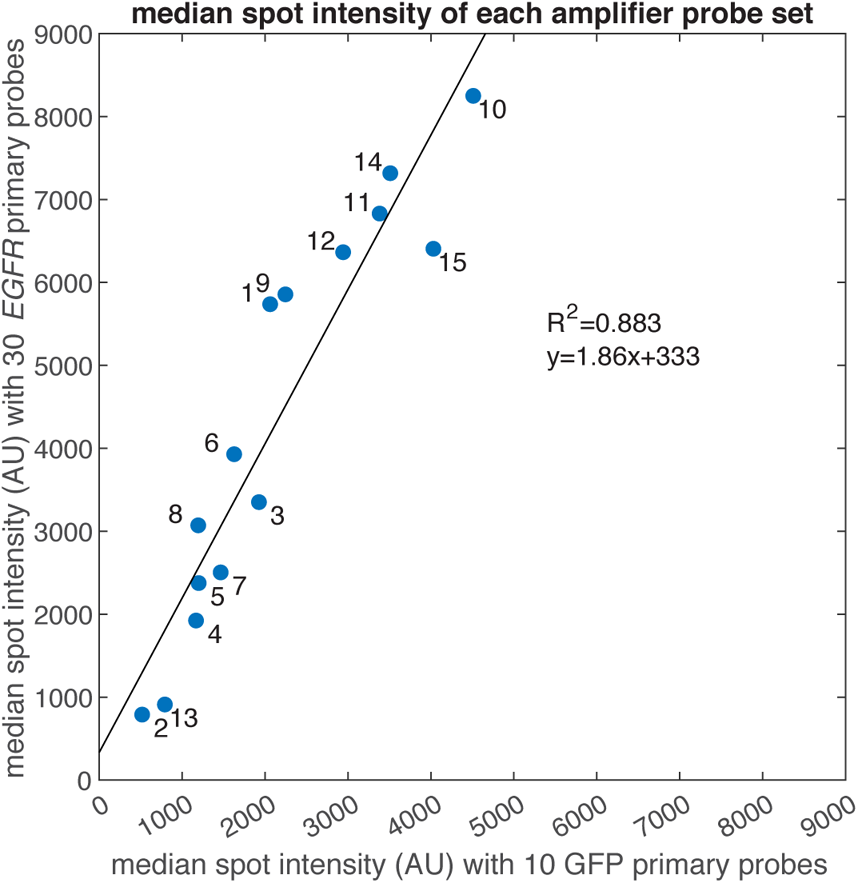
Amplifier probe sets can modularly be used with various primary probe sets. We plotted the median spot intensity generated by each clampFISH 2.0 amplifier set (labeled 1 to 15) from the amplifier screen experiment when used with primaries targeting GFP (x-axis) or *EGFR* (y-axis). We observed a strong correlation (R^2^=0.883) between the two primary probe sets, suggesting that gene-specific effects on amplification play a minimal role in their performance. The slope of the regression suggests a nearly 2-fold increase in spot intensities when amplifier sets were used with the *EGFR* probe set over the GFP probe set, likely as a result of the 3-fold higher number of primary probes (30 for *EGFR* vs. 10 for GFP).

**Supplementary Figure 7:**
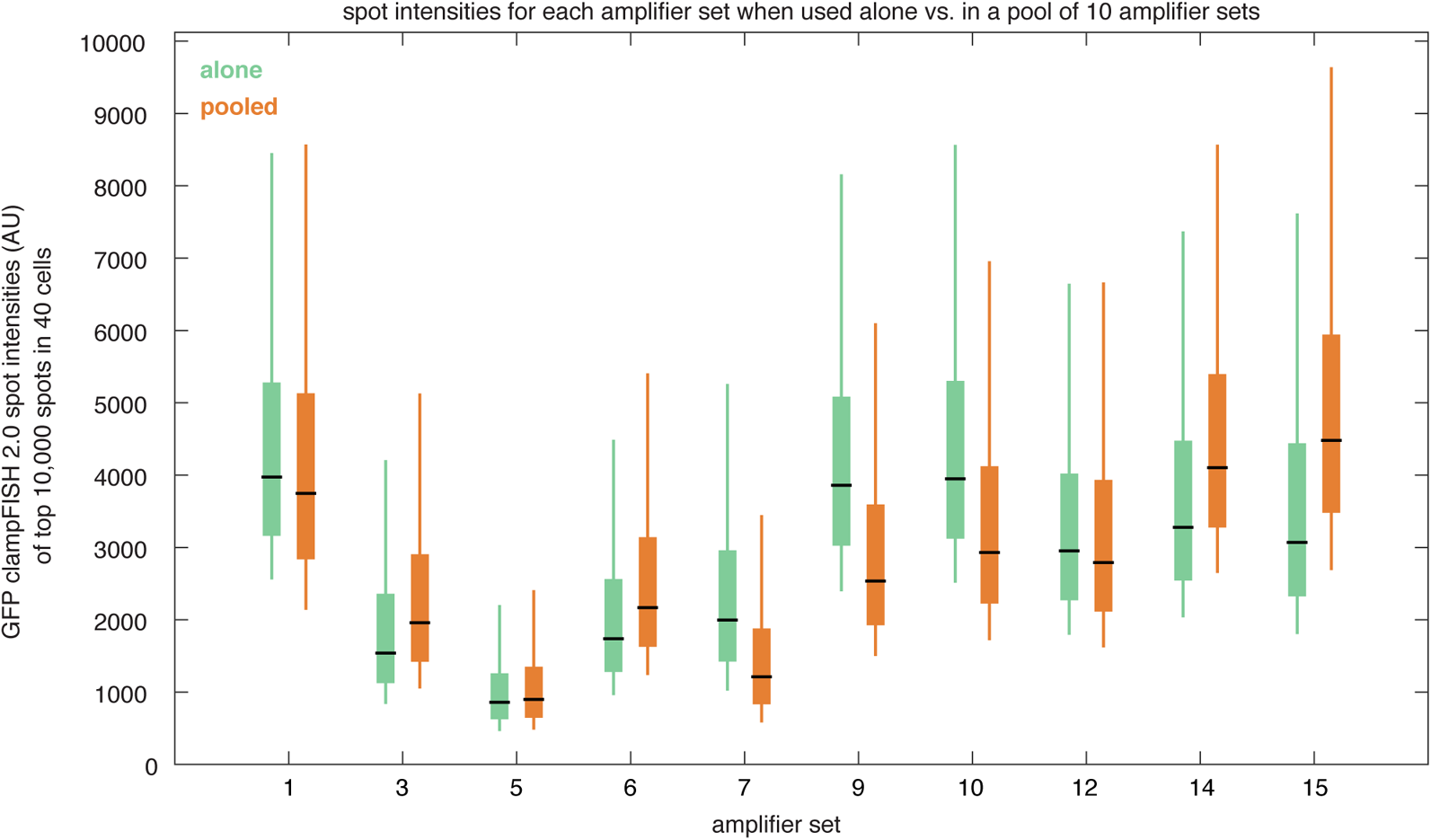
Amplifier probe sets amplify signal similarly when used alone vs. when used in a pool of 10 amplifier probes. We hybridized 10 GFP-targeting primary probe sets, with each set ligated to a different amplifier-binding oligonucleotide, and amplified each in one of two ways: with its corresponding amplifier probe set alone (green) or with a pool of all 10 amplifier sets (orange). Plotted are the intensities of the 10,000 highest-intensity spots from 40 segmented cells per condition (379 ‘alone’ spots outliers and 418 ‘pooled’ spot outliers not shown).

**Supplementary Figure 8:**
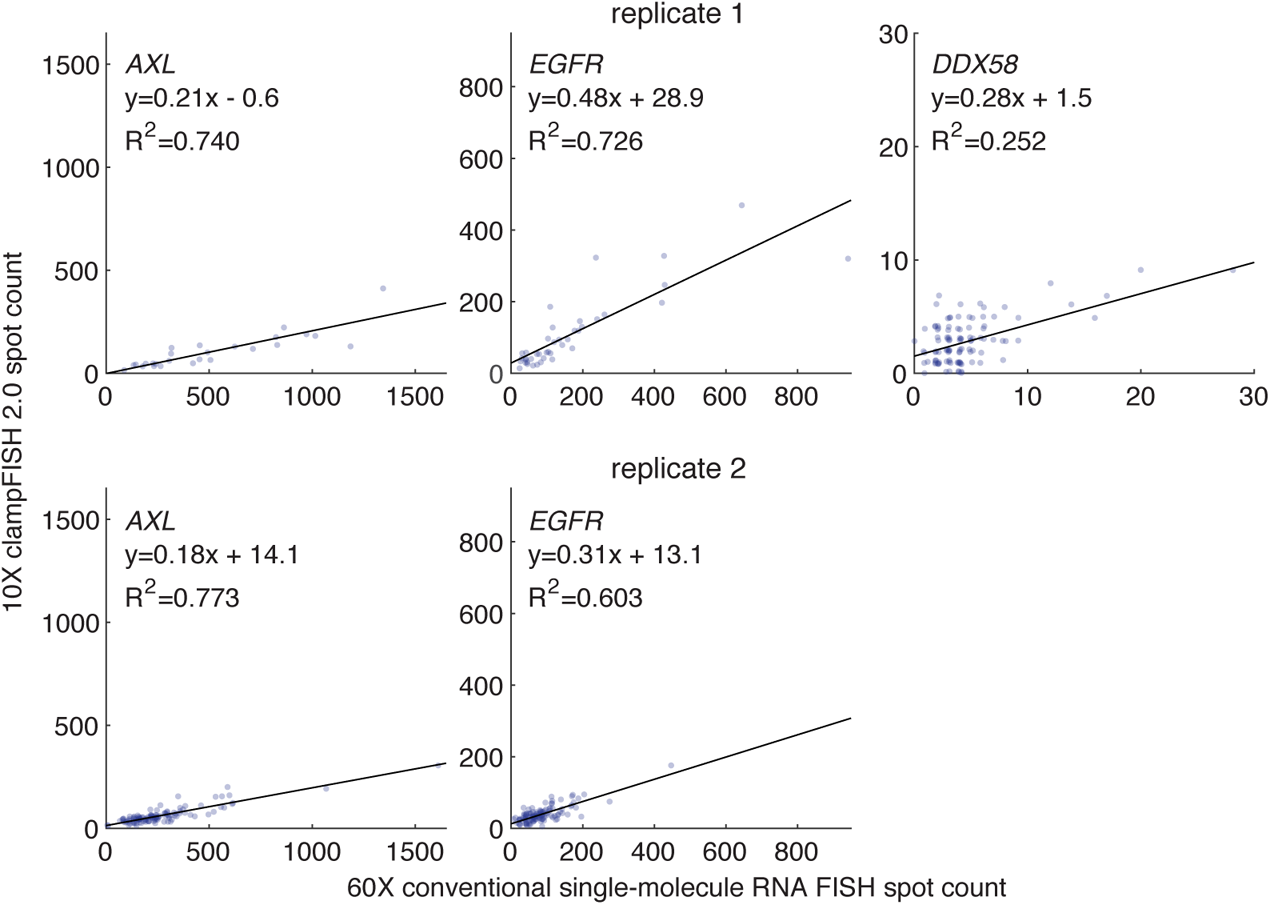
clampFISH 2.0 quantifies RNA spot counts at 10X magnification. Depicting the same data as in Fig. 2b, but with clampFISH 2.0 spots imaged at 10X magnification. We performed clampFISH 2.0 for 10 genes, amplified the 10 scaffolds in parallel to round 8, then added a single pair of readout probes to label a scaffold corresponding to *AXL* (left; in drug-resistant WM989 A6-G3 RC4 cells), *EGFR* (middle; in drug-resistant WM989 A6-G3 RC4 cells), or *DDX58* (right; in drug-naive WM989 A6-G3 cells). In two biological replicates (top: replicate 1; bottom: replicate 2), we counted spots for clampFISH 2.0 at 10X magnification (y-axis) and conventional single-molecule RNA FISH at 60X magnification (x-axis), which targeted non-overlapping regions of the same RNAs. In replicate 2, imaging at 10X of *DDX58* spots before conventional single-molecule RNA FISH was not performed.

**Supplementary Figure 9:**
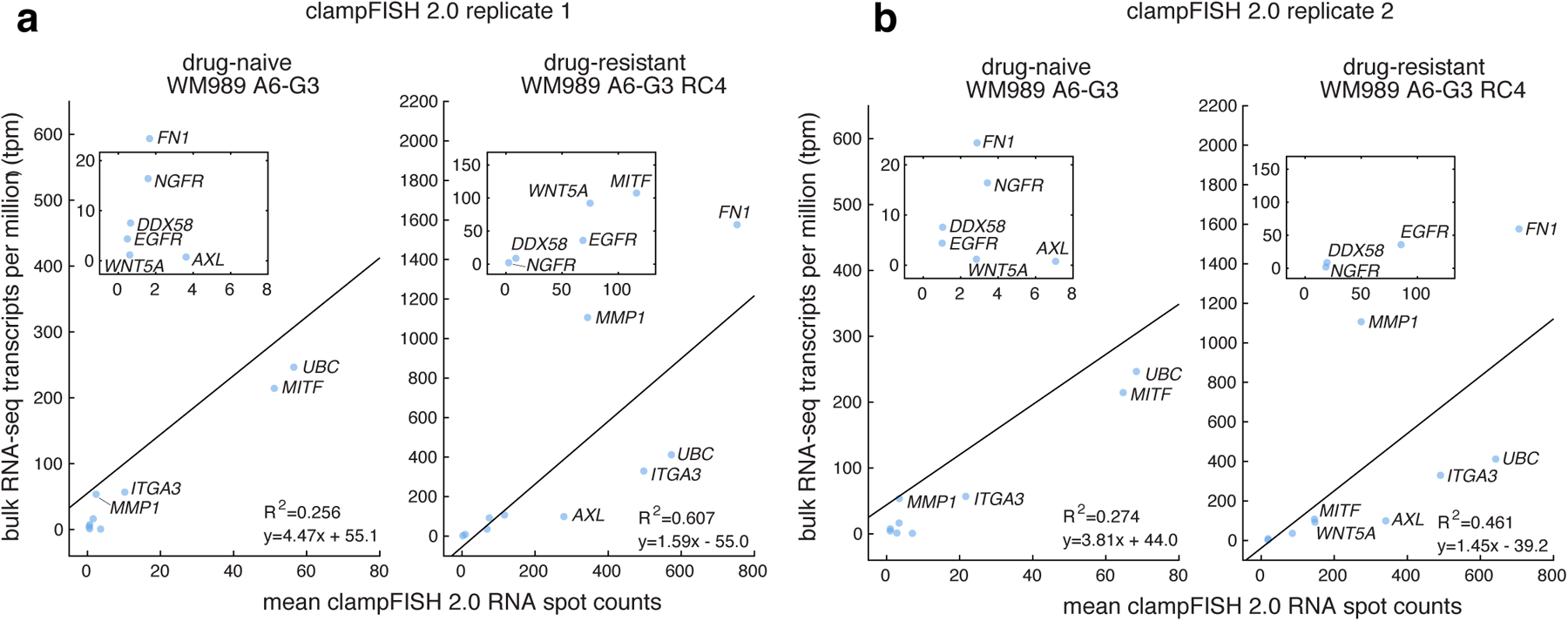
Mean clampFISH 2.0 spot counts are correlated with bulk RNA sequencing data. **(a)** Mean clampFISH 2.0 spot counts from 722,298 drug-naive WM989 A6-G3 cells (left) and 2,155 vemurafenib-resistant WM989 A6-G3 RC4 cells (right) for the 10 genes from the high-throughput profiling experiment (x-axis) and bulk RNA-seq transcripts per million (y-axis) for each of the two cell lines. **(b)** A technical replicate of the same experiment as in (a), but with data from 234,410 drug-naive WM989 A6-G3 cells and 5,150 vemurafenib-resistant WM989 A6-G3 RC4 cells, using the same bulk RNA sequencing data. We observed that *FN1* and *MMP1*, both of which have a lower mean clampFISH 2.0 spot count than would be expected from the remaining genes’ trend, are expressed at particularly high levels in a subset of cells (see Fig. 3b), suggesting that optical crowding at 20X magnification may contribute to their under-counting by clampFISH 2.0.

**Supplementary Figure 10:**
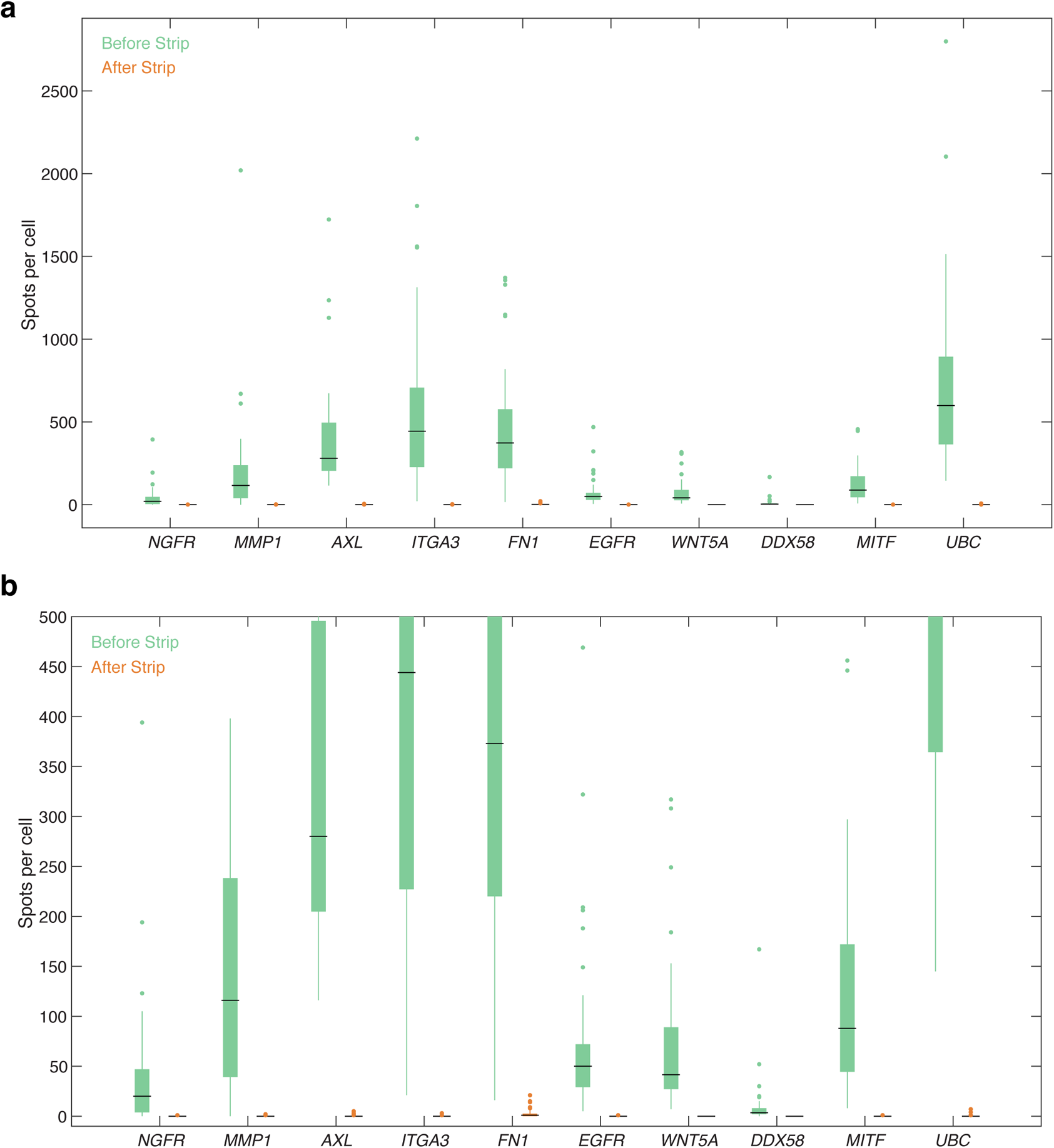
clampFISH 2.0 readout probe signal can be removed with a high-stringency wash. **(a)** Boxplots of clampFISH 2.0 spots per cell detected above a chosen gene-specific threshold for 10 genes before (green) and after (orange) the readout probe stripping protocol. Shown for each gene are spot counts from one of two melanoma lines with higher expression for that gene (for *NGFR*: drug-naive WM989 A6-G3 cells; for all other genes: vemurafenib-resistant WM989 A6-G3 RC4 cells). Each condition contains 39-48 segmented cells where each cell is represented in both the before-stripping and the after-stripping data. The box and whiskers for the after strip data are at 0 spots and thus are not visible, except for *FN1* which has an interquartile range from 0 to 2.5 spots and a whisker extending to 6 spots. **(b)** Depicting the same data as in (a) for only data below 500 spots per cell.

**Supplementary Figure 11:**
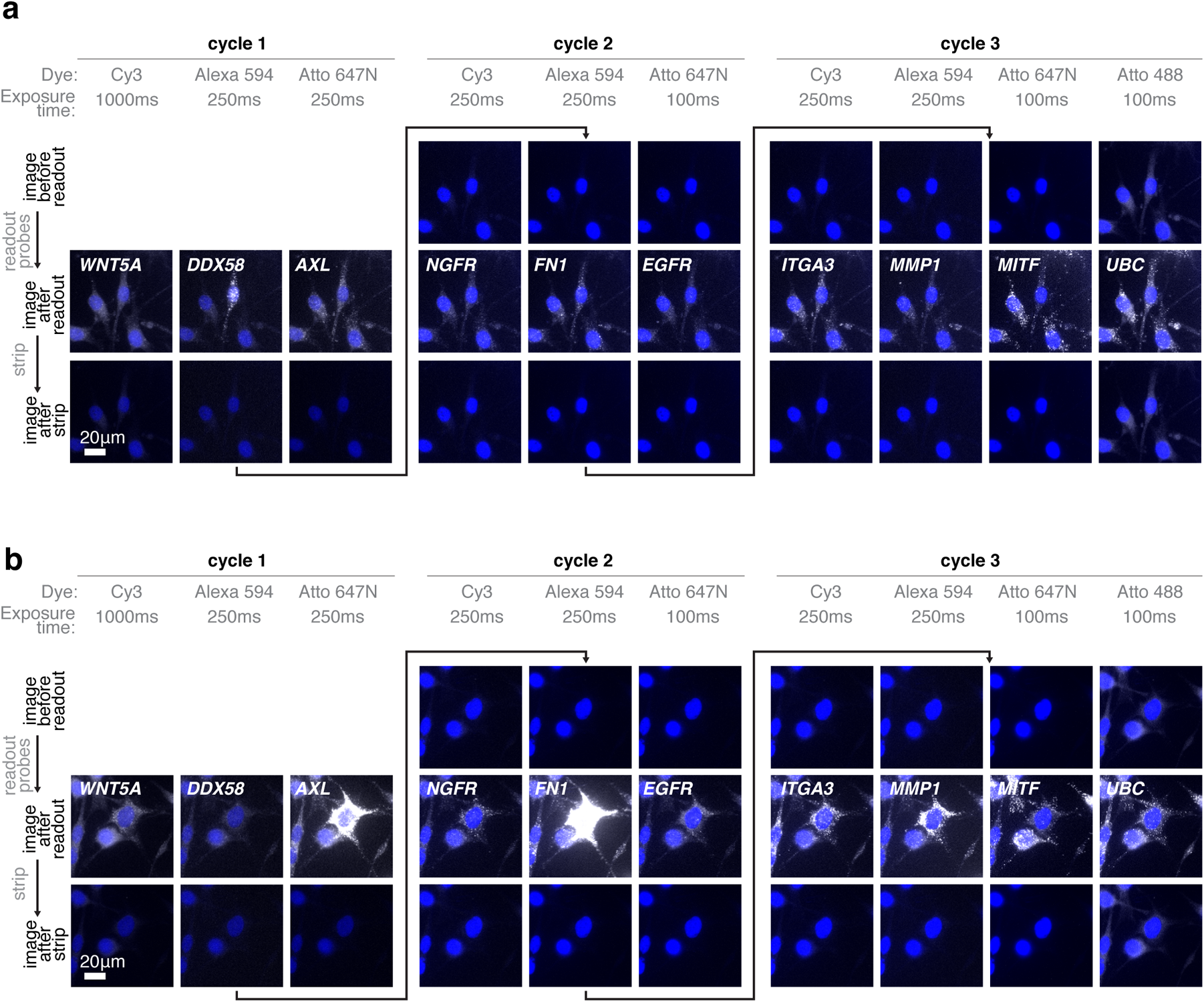
Signal from the previous readout cycle is removed after a high-formamide strip. **(a)** Example images of clampFISH 2.0 spots at 20X magnification before the readout probe hybridization (top row), after adding readout probes (middle row), and after stripping off readout probes (bottom row). The first three columns are from readout cycle 1, the next three are from readout cycle 2, and the last 4 columns are from readout cycle 3. Each column’s images are from the same channel (with the corresponding readout probe dye indicated), exposure time (as indicated in milliseconds), and are contrasted identically. **(b)** Example images as in (a) at a different position on the plate.

**Supplementary Figure 12:**
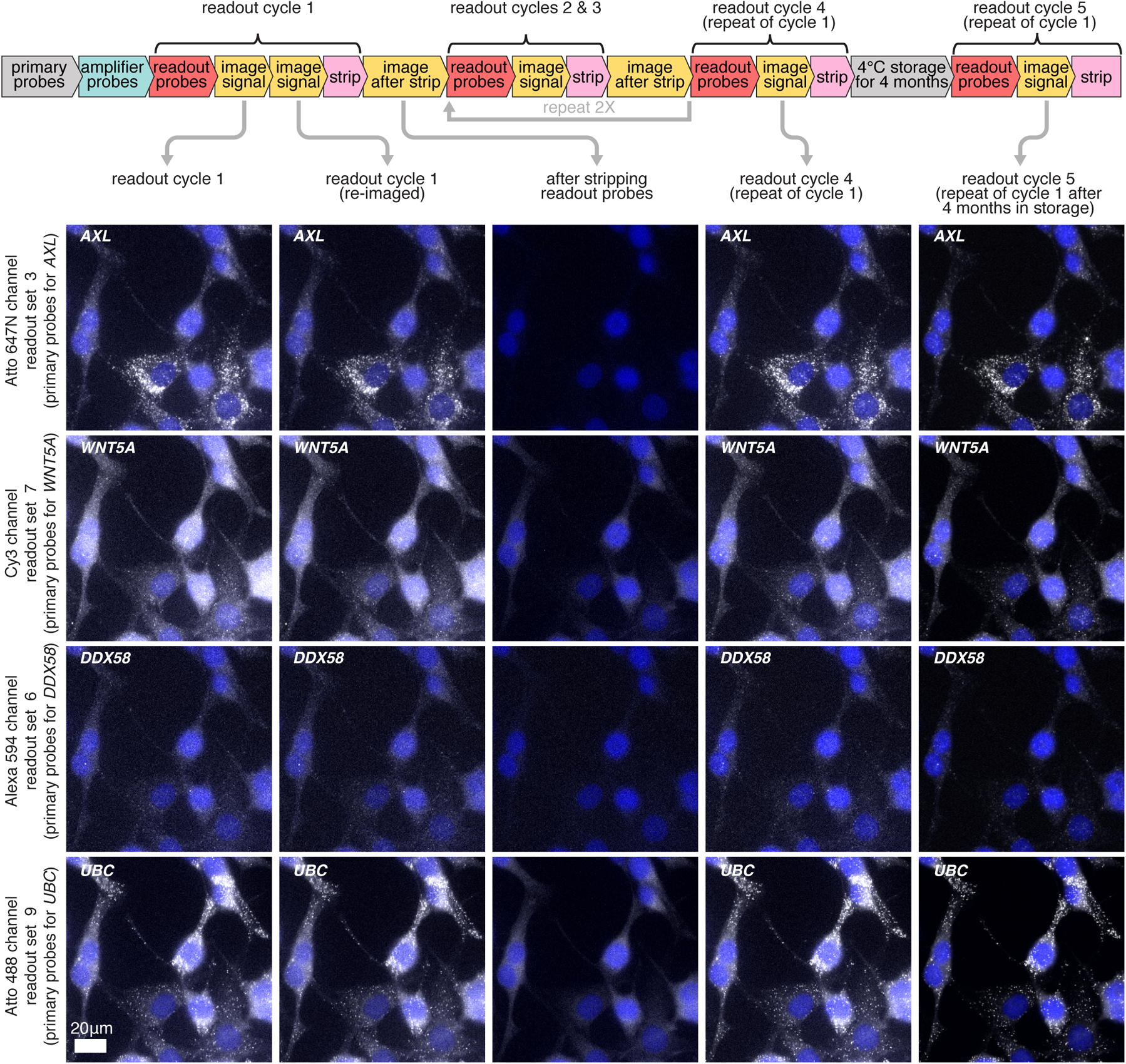
clampFISH 2.0 scaffolds remain stably bound after multiple rounds of readout stripping and storage at 4°C for 4 months. Images of clampFISH 2.0 spots from a 20X objective over readout cycles where we repeatedly use 4 sets of readout probes which label (from top to bottom) *AXL*, *WNT5A*, *DDX58*, and *UBC* clampFISH 2.0 scaffolds. **Column 1**: readout cycle 1. **Column 2**: readout cycle 1, re-imaged after removing the sample from the microscope stage and stored overnight at 4°C. **Column 3**: after stripping off readout probes from readout cycle 1. **Column 4**: readout cycle 4, where we repeat readout cycle 1 after readout cycles 2 and 3 (where different sets of genes were labeled). **Column 5**: readout cycle 5, performed after storing the sample at 4°C in 2X SSC for 4 months. DAPI overlay is contrasted separately for each column. Each row of readout cycle 5 (column 5) is contrasted with 180% the intensity range of the first four columns. The cycle 5 signal presumably appeared brighter due to changes in the microscope’s optics during that time frame (eg. greater sample illumination or increased transmission to the sensor).

**Supplementary Figure 13:**
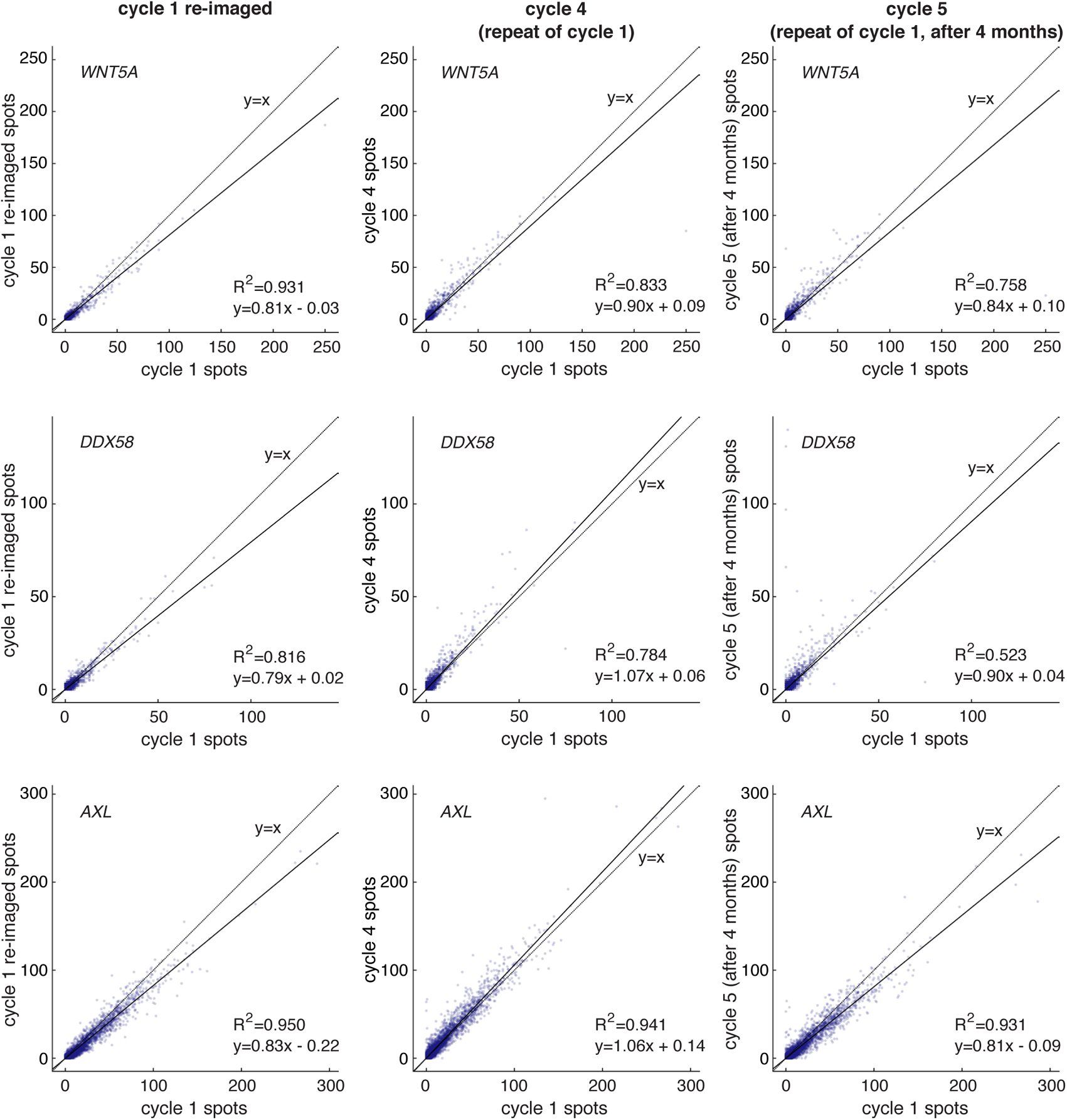
clampFISH 2.0 scaffolds remain stably bound after multiple rounds of readout stripping (replicate 1) and storage at 4°C for 4 months. clampFISH 2.0 spots per cell for (from top to bottom) *WNT5A*, *DDX58*, and *AXL* from readout cycle 1 (x-axis) plotted against 3 additional rounds of imaging for the same probed scaffold: re-imaged readout cycle 1 (column 1 plots); readout cycle 4, where the same readout probes were used as cycle 1 (column 2 plots); and readout cycle 5, where again we used the same readout probes as cycle 1 after being stored for 4 months in 4°C (column 3 plots). Each point is one of 44,227 cells. See Supplementary Figure 12 for experiment workflow schematic. Shown is one of two technical replicates (see Supplementary Figure 14 for replicate 2).

**Supplementary Figure 14:**
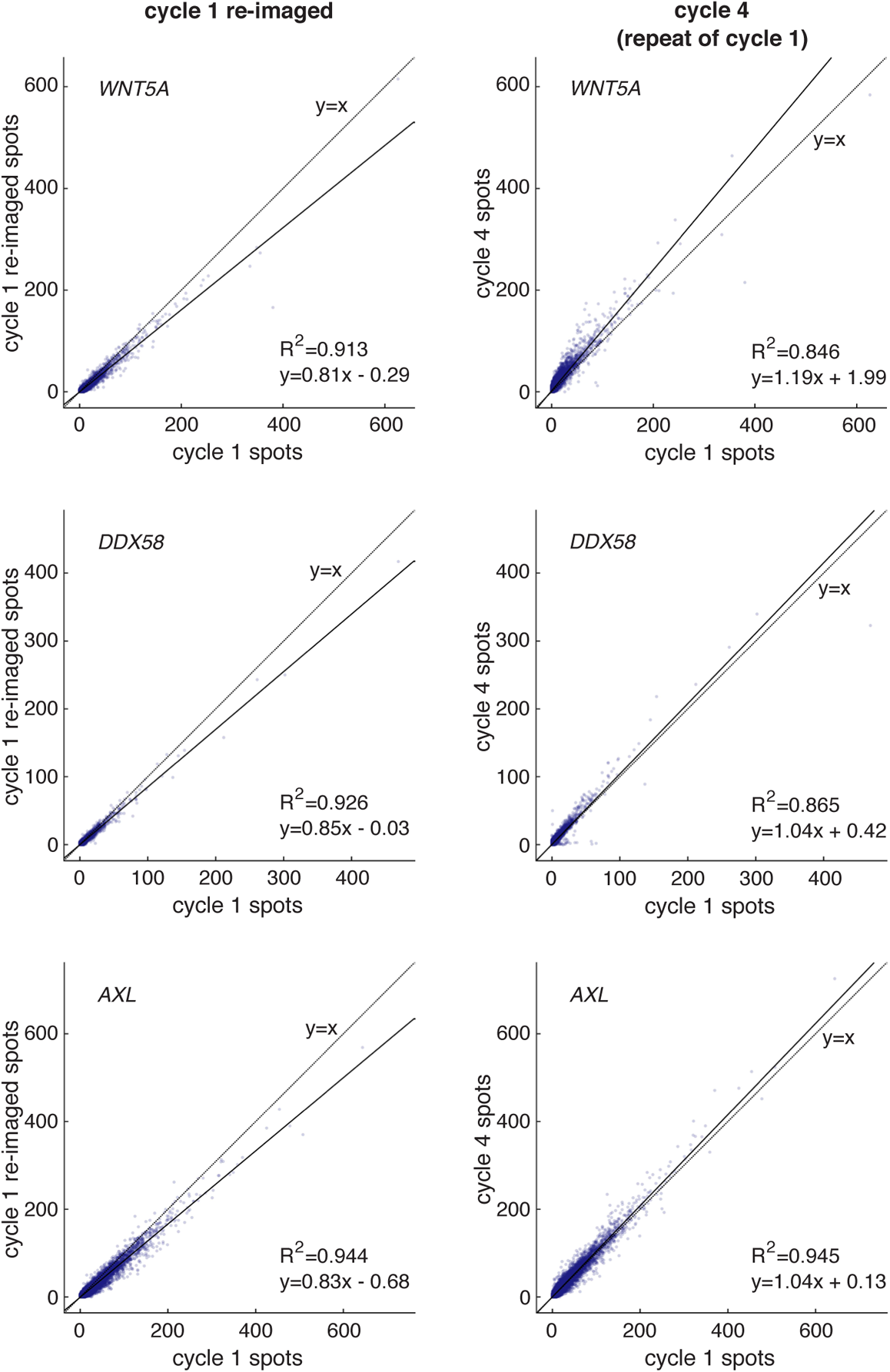
clampFISH 2.0 scaffolds remain stably bound after multiple rounds of readout stripping (replicate 2). Technical replicate 2 of the experiment from Supplementary Figure 13, but without readout cycle 5. clampFISH 2.0 spots per cell for (from top to bottom) *WNT5A*, *DDX58*, and *AXL* from readout cycle 1 (x-axis) plotted against 2 additional rounds of imaging for the same probed scaffold: re-imaged readout cycle 1 (column 1 plots); and readout cycle 4, where the same readout probes were used as cycle 1 (column 2 plots). Each spot is one of 89,545 cells. See Supplementary Figure 12 for experiment workflow schematic.

**Supplementary Figure 15:**
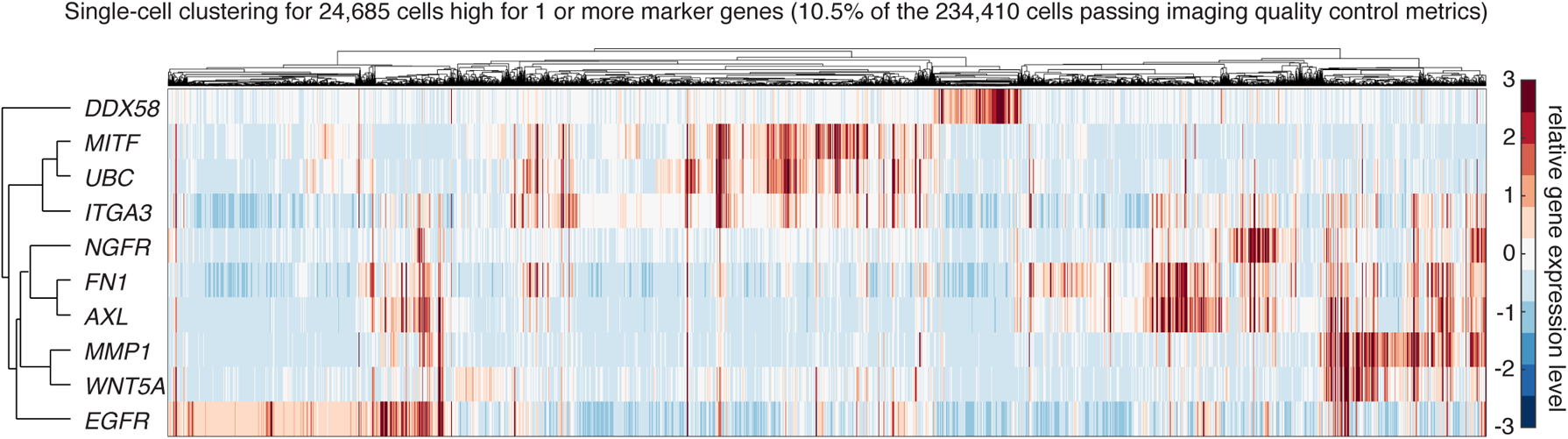
Clustering of cells expressing one or more drug resistance markers. Technical replicate 2 of the high-throughput profiling experiment from Fig. 3c. clampFISH 2.0 was performed for 10 genes in 253,662 drug-naive WM989 A6-G3 cells. We detected 24,685 cells (10.5% of the 234,410 cells passing quality control checks) that had high levels of one or more of 8 cancer marker genes (*WNT5A*, *DDX58*, *AXL*, *NGFR*, *FN1*, *EGFR*, *ITGA3*, *MMP1*) and performed hierarchical clustering on this population. See methods section for cutoff values defining high expression levels.

**Supplementary Figure 16:**
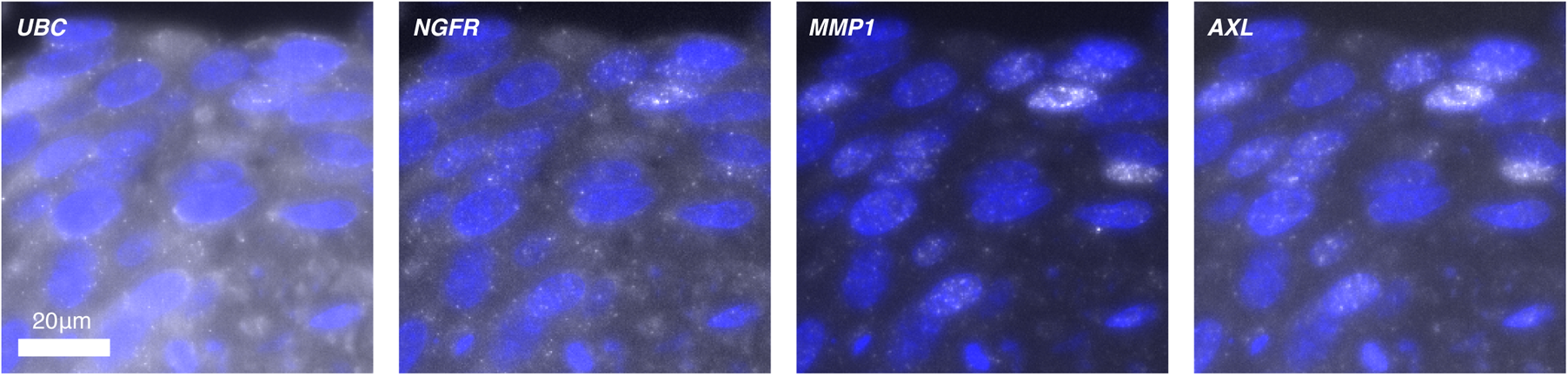
clampFISH 2.0 in FFPE tissue. We performed clampFISH 2.0 for ten genes in formalin-fixed paraffin embedded (FFPE) tumor tissue derived from human WM4505-1 cells injected into a mouse. We then hybridized readout probes for four clampFISH 2.0 scaffolds, from left to right: *UBC* (Atto 488), *NGFR* (Cy3), *MMP1* (Alexa Fluor 594), and *AXL* (Atto 647N). Shown are images that were taken at 20X magnification.

**Supplementary Figure 17:**
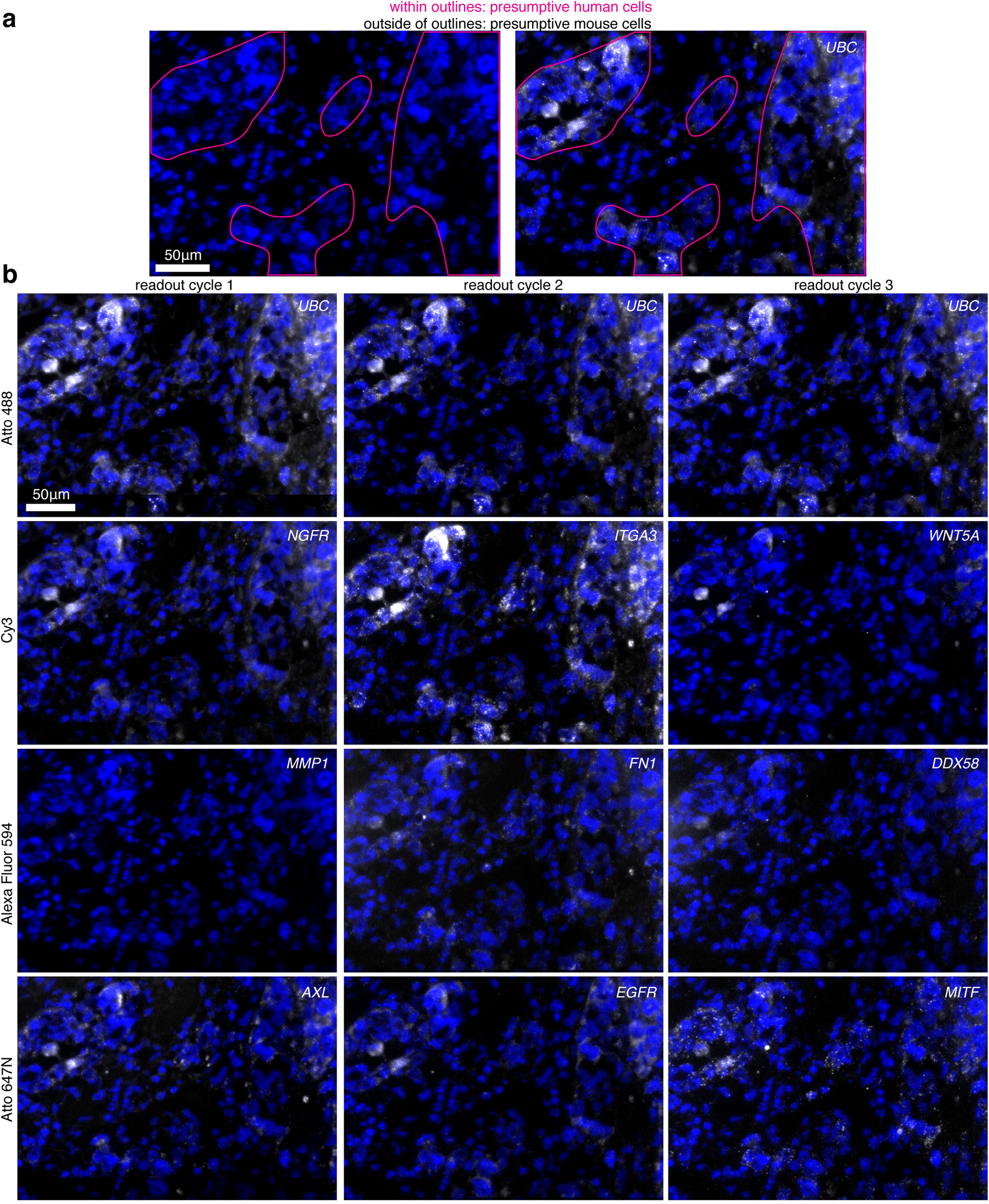
clampFISH 2.0 detects RNAs in presumptive human cells in tissue. clampFISH 2.0 was performed in a 6μm fresh frozen tissue section of a dissected tumor, derived from human WM989-A6-G3-Cas9-5a3 cells injected into a mouse and fed chow containing the BRAF^V600E^ inhibitor PLX4720. Shown are stitched maximum intensity projections of 20X image stacks with 5 z-planes at 1.2μm z-step increments. **(a)** Pink outlines around regions containing mostly presumptive human cells, demarcated based on nuclear morphology, showing DAPI staining alone (left) and DAPI with UBC clampFISH 2.0 signal overlaid (right), where images are from readout cycle 2. **(b)** clampFISH 2.0 scaffolds for 10 genes were probed across readout cycles 1 (left), 2 (middle), and 3 (right), where the UBC scaffold was probed each round as a positive control. The dyes on each readout probe set were (top to bottom): Atto488, Cy3, Alexa Fluor 594, and Atto647N.

**Supplementary Figure 18:**
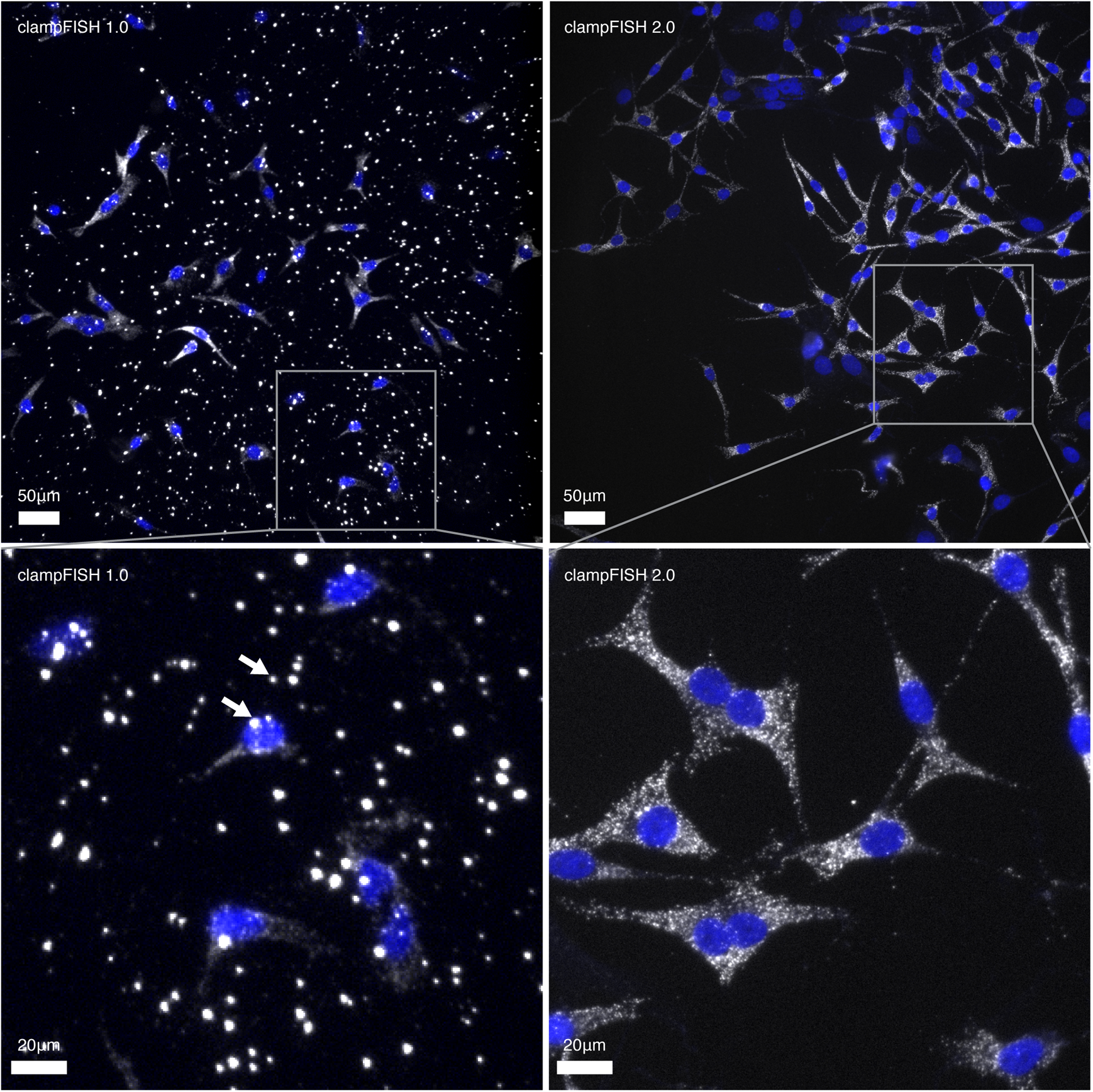
clampFISH 2.0 eliminates the bright, non-specific fluorescent spots that were observed in clampFISH 1.0. **Top left**: clampFISH 1.0 targeting GFP in WM983b-GFP melanoma cells, amplified to round 6 with amplifier probes containing an internal Cy5 dye and imaged at 20X with a 3 second exposure time using a cooled CCD camera with a 13μm pixel size (image from Rouhanifard et al. 2018; see that paper for further details). The two arrows point to two of the non-specific spots. **Top right**: clampFISH 2.0 targeting GFP in a mixed population of cells (a majority of WM989 A6-G3 H2B-GFP cells and fewer WM989 A6-G3 RC4 cells), amplified to round 8 with readout probes labeled with Atto 647N and imaged at 20X with a 1 second exposure time using a sCMOS camera with a 6.5μm pixel size (image from the present work’s pooled amplifier experiment). **Bottom**: zoomed-in views of the top images. We found that we could eliminate the bright non-specific spots by introducing a number of centrifugation steps to both the primary probe and amplifier probe synthesis protocols. To perform this step, we centrifuge the solution in 1.5mL tubes at 17,000g for 20 minutes and transfer the top portion of the solution to a new tube and discarded the bottom portion. We perform this step twice after the enzymatic steps are complete, and once after ethanol precipitation. Additionally, we found that by adding the centrifugation step to completed clampFISH 1.0 probe solutions, we could similarly reduce the non-specific spots seen in that method.

